# Pairing Metagenomics and Metaproteomics to Characterize Ecological Niches and Metabolic Essentiality of gut microbiomes

**DOI:** 10.1101/2022.11.04.515228

**Authors:** Tong Wang, Leyuan Li, Daniel Figeys, Yang-Yu Liu

## Abstract

The genome of a microorganism encodes its potential functions that can be implemented through expressed proteins. It remains elusive how a protein’s selective expression depends on its metabolic essentiality to microbial growth or its ability to claim resources as ecological niches. To reveal a protein’s metabolic or ecological role, we developed a computational pipeline, which pairs metagenomics and metaproteomics data to quantify each protein’s gene-level and protein-level functional redundancy simultaneously. We first illustrated the idea behind the pipeline using simulated data of a consumer-resource model. We then validated it using real data from human and mouse gut microbiome samples. In particular, we analyzed ABC-type transporters and ribosomal proteins, confirming that the metabolic and ecological roles predicted by our pipeline agree well with prior knowledge. Finally, we performed *in vitro* cultures of a human gut microbiome sample and investigated how oversupplying various sugars involved in ecological niches influences the community structure and protein abundance. The presented results demonstrate the performance of our pipeline in identifying proteins’ metabolic and ecological roles, as well as its potential to help us design nutrient interventions to modulate the human microbiome.

## Introduction

Metagenomic sequencing has enabled the measurement of the genomic content and functional potential of microbial communities at an unprecedented rate, aiding the understanding of their role in host health^1–3^ and biogeochemical cycling^4–6^. While various computational approaches based on these genomes quantify interactions within microbial communities^7–12^ and analyze the functional redundancy and functional stability of microbial communities^10–12^, they focus on potential rather than actual function, as microorganisms only express a subset of genes as proteins^13^. Recent advancements in high-throughput metaproteomics allow us to quantify protein abundances in human gut microbiomes^14^, offering insights into gene expression in response to environmental changes when paired with metagenomic data.

From the metabolic perspective, some genes and their encoded proteins are indispensable for cell metabolism under any conditions, and microbial growth halts without these essential functions—aminoacyl-tRNA synthetase^15,16^, ribosomal proteins^17–19^, and enzymes involved in glycolysis^20,21^. From the ecological perspective, gene expression is influenced by ecological selection, with specific proteins indicating which resources a microbe can utilize and defining its ecological niche. For instance, *E. coli* prefers glucose over lactose due to the repressed expression of lactose-utilizing enzymes, even though it can use both sugars^22,23^. Such specialization of consuming one resource caused by the selective gene expression may reduce the niche overlap with other species and allow microbial coexistence, as seen with two *E. coli* strains where one expresses acetyl-coenzyme synthetase (Acs)^24,25^ to consume acetate produced by the other^26–29^.

Understanding the selective expression of microbial genes is an outstanding question in microbiology. Does the behavior of selective expression of microbial genes differ between metabolic function (e.g., essential for microbial growth metabolism) and ecological function (e.g., claiming resources as a niche)? To answer this, we developed a computational method to analyze paired metagenomic and metaproteomic^30–33,14^ data, constructing the gene content network (GCN) or protein content network (PCN) --- a bipartite graph that connects microbial taxa to their genes or expressed proteins, respectively (Figs. 1a&b). For each gene and its encoding protein, we compared its gene-level (or protein-level) functional redundancy (FR), revealing each protein family’s metabolic or ecological role. Our method, validated with several gut microbiome data, accurately predicts that ABC-type transporters are related to ecological niches^34–36^, and ribosomal proteins are essential^17–19^. Finally, we performed *in vitro* culture experiments using human gut microbiome samples to investigate how oversupplying sugars involved in ecological niches influence community structure and protein expression.

**Figure 1:**
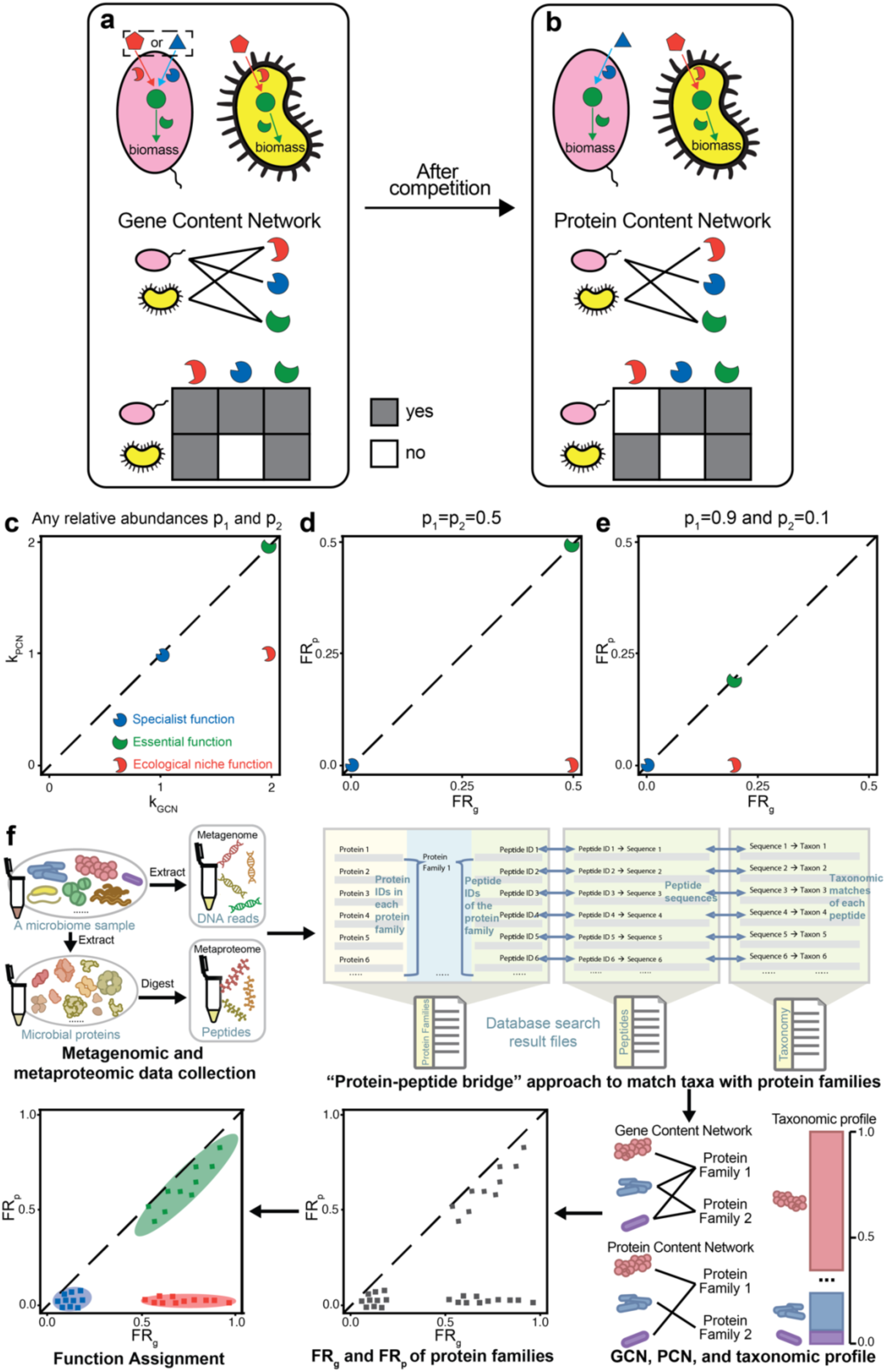
Protein functions involved in determining ecological niches are postulated to have larger discrepancies between the gene-level functional redundancy FR_g_ and protein-level functional redundancy FR_p_. Here we use a hypothetical example with three representative proteins (3 broken circles with complementary shapes to their substrates) to demonstrate this point. a, Schematic of the genomic capacity of two microbial taxa (pink oval vs yellow indented oval). Two resources (red pentagon and blue triangle) are externally supplied to the community. The green metabolite can be transformed from the red or blue resource and further utilized in biomass synthesis. The pink taxon has the capacity of converting either supplied resource into the green metabolite (red and blue arrows), while the yellow taxon can only convert the red resource (red arrow). b, Schematic of expressed proteins for two microbial taxa after their competition in the same community. After the competition, the reduced resource conflict (represented by the pink taxon choosing the blue resource as the sole one to consume) can promote their coexistence. Gene content network (GCN) and protein content network (PCN) can be used to capture genomic capacity and expressed protein functions for all taxa. Alternatively, this network can be represented as incidence matrices on the bottom (grey areas imply the existence of edges connecting taxa to proteins). c-d, The comparison between k_GCN_ and k_PCN_ or between FR_g_ and FR_p_ helps to classify proteins into three protein functional types: specialist function, essential function, and niche function. In the calculation of FR_g_ and FR_p_, we assume equal abundances of the two species, i.e., *p*_1_ = *p*_2_ = 0.5. e, The comparison between FR_g_ and FR_p_ when *p*_1_ = 0.9 and *p*_2_ = 0.1. f, The pipeline of assigning the functions (specialist, niche, or essential) to protein families based on the paired metagenomes and metaproteomes. Each individual’s gut microbiome sample was subjected to DNA and protein extraction. Then a protein-peptide bridge approach can be used for generating the GCN based on the metagenome and PCN based on the metaproteome. When matched metagenomes are available, taxonomic and functional annotations of the metagenomes can be used for PCN generation. Based on the generated GCN, PCN, and taxonomic profile, FR_g_ and FR_p_ can be computed and used for the function assignment.

## Materials and Methods

### *In-vitro* human gut microbiota culture and metaproteomics

Three healthy individual microbiota samples were collected and biobanked. The frozen microbiome samples were cultured in our optimized culture medium^37^ with or without the presence of different sugars in technical triplicates, and were taken at different times for optical density and metaproteomic analyses. For single-strain samples, proteins were extracted with 4% SDS 8M urea buffer in 100 mM Tris-HCl buffer, followed by precipitation and acetone washing. Proteins were digested with trypsin desalted^38^ for LC-MS/MS analysis using an Orbitrap Exploris 480 mass spectrometer. For the cultured microbiomes, an automated process extracted and purified proteins, which were then digested, desalted, and quantified using TMT11plex^39^, ensuring mixed representation in labeling to avoid bias. Samples underwent a 2-hour LC gradient and were analyzed by mass spectrometry. More details can be found in Supplemental Methods.

### Datasets

Metagenomics data from four individual microbiomes were obtained from the previous MetaPro-IQ study^14,33^ (accessible from the NCBI sequence read archive (SRA) under the accession of SRP068619) and the same samples were reanalyzed by an ultra-deep metaproteomics approach^14^ via the PRIDE partner repository^40^ with the dataset identifier PXD027297). Proteomics dataset of the cultured singles strain samples has been deposited to ProteomeXchange Consortium with the identifier PXD037923. Metaproteomic dataset of the RapidAIM-cultured microbiome samples has been deposited to ProteomeXchange Consortium (identifier PXD037925). The metaproteomic dataset of the mouse gut microbiome comprising twenty gut microbes is derived from a previous study^41^ that was deposited to ProteomeXchange Consortium with the dataset identifier PXD009535 and to MassIVE with the dataset identifier MSV000082287.

### Database search and data processing

Proteomics database searches used FASTA databases of the individual strains downloaded from NCBI and MaxQuant^42^ 1.6.17.0 for analysis, without the label-free quantification. Metaproteomic database searches of cultured microbiome samples were performed using MetaLab V2.2^43^, MaxQuant option was used to search the TMT dataset against the IGC database of the human gut microbiome. The resulting data table was normalized using R package MSstatsTMT, and missing values were imputed using R package DreamAI^45^. The “fraction” of each taxon-specific protein is computed by dividing the protein intensity by the sum of the intensities of all proteins assigned to the same taxon. The log2 fold change of each protein is obtained by taking log2 of the ratio between its fraction in the treatment group (with added sugars) and its fraction in the control group (without added sugars).

### Statistics

To calculate correlation throughout the study, we used Pearson’s correlation coefficient. All statistical tests were performed using standard numerical and scientific computing libraries in the Python programming language (version 3.7.1) and Jupyter Notebook (version 6.1).

## Results

### Specialist function, niche function, and essential function

Here, we define three types of functions for protein families that we would like to categorize: (1) “*Specialist function*”: specialized by only a few taxa and not widely shared within a community. (2) “*Niche function*”: arising from ecological competition, widespread among genomes of numerous taxa but selectively expressed under specific ecological conditions. (3) “*Essential function*”: metabolically indispensable for and widely shared by many taxa within a microbial community. We emphasize that our definition is not exhaustive; some proteins may display attributes of multiple categories or not align precisely with any single category.

Using a simple hypothetical example of two competing species (Fig. 1a, b), we demonstrated the three function types: (1) The blue protein is a specialist function since it is solely encoded in the pink species’ genome; (2)The red protein belongs to a niche function due to its selective expression by the yellow species even though the protein is encoded in the genomes of both species; (3) The green protein is an essential function because both species need it for biomass synthesis. In coexistence, the pink species specializes in the blue resource, avoiding competition with the yellow species for the red resource.

### GCN, PCN, and network degree

We can identify the functional types of proteins in this hypothetical case by comparing the structure of the GCN and PCN (Fig. 1a, b). For example, consider the protein responsible for converting red resource to green metabolite (red broken circle in Fig. 1a, b), its degree in the GCN k_GCN_ = 2, while its degree in the PCN k_PCN_ = 1. This degree reduction is due to distinct ecological niches being occupied by two species when they are cocultured. By contrast, the protein responsible for assimilating critical green metabolites (green broken circle in Fig. 1a, b) into biomass does not show a degree reduction (k_GCN_ = k_PCN_ = 2) because it is essential for microbial growth. Similarly, since the blue protein is only specialized by the pink species, its k_GCN_ = k_PCN_ = 1. Thus, three function types occupy different regions in the k_GCN_ vs. k_PCN_ plot (Fig. 1c).

### Quantifying gene- and protein-level functional redundancy of each gene and its encoded protein

However, the network degree does not consider the significant impact of the microbial taxonomic profile, which details the makeup of a microbial community. This profile is represented by ***p*** = (*p*_1_, …, *p*_*N*_), where *p*_*i*_ is the relative abundance of taxon-*i* and 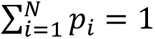. For a given gene and its encoded protein, we can define its gene-level FR (FR_g_) and protein-level FR (FR_p_) within this sample as

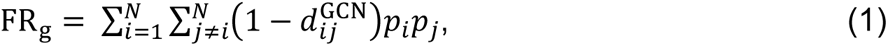

and

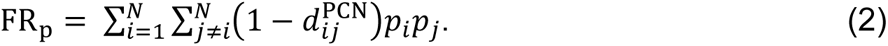

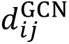 (or 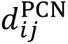) is the distance between taxon-*i* and taxon-*j* based on their genomic capacity to express this gene (or the presence of the protein). For simplicity, we assume 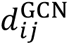 is binary, i.e., 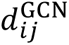 = 0 if and only if both taxa share the potential to express the gene, and 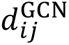 = 1 otherwise. 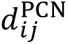 = 0 if and only if both taxa have expressed the protein. Here, we define FR_g_ and FR_p_ for each protein, different from our previous studies where FR was calculated by including all genes or proteins in a microbial community^12,14^.

Comparing FR_g_ and FR_p_ provides deeper insight into proteins’ function types. For the red protein in our hypothetical example, 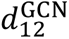 = 0 and 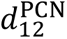 = 1 because both species share the potential to express the gene while only the yellow species have expressed it (Fig. 1a, b). As a result, FR_g_ = 2(1 − 0)*p*_1_*p*_2_ = 2*p*_1_*p*_2_ and FR_p_ = 2(1 − 1)*p*_1_*p*_2_ = 0 (Fig. 1d, e). Following the same analysis, FR_g_ = FR_p_ = 2*p*_1_*p*_2_ for the green protein, and FR_g_ = FR_p_ = 0 for the blue protein (Fig. 1d, e). Different from composition-independent k_GCN_ and k_PCN_, FR_g_ and FR_p_ take the microbial composition into account and thus are more ecologically meaningful. Notably, a more uneven abundance distribution would lead to smaller FR_g_ and FR_p_ (*p*_1_ = *p*_2_ = 0.5 in Fig. 1d; *p*_1_ = 0.9 and *p*_2_ = 0.1 in Fig. 1e). The influence of relative abundances on FR can be mitigated by using the normalized FR: nFR = FR / TD, where 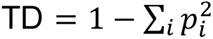 (Supplementary Fig. 1; see Supplementary Information for the definition).

#### Overview of our computational pipeline

Following the idea of comparing FR_g_ with FR_p_, we developed a computational pipeline to assign the function types (specialist, niche, or essential) to protein families based on the paired metagenome and metaproteome (Fig. 1f). This pipeline starts with DNA sequences from metagenomes and peptides sequences from metaproteomes. Using the "protein-peptide bridge" approach that maps peptides to their taxonomic origins and protein families (i.e., orthologous protein clusters), it generates the GCN, PCN, and taxonomic profile, from which we compute FR_g_ and FR_p_. Details about this approach can be found in the Supplementary Information. Finally, based on the scatterplot of FR_g_ vs. FR_p_, each protein family is categorized into one of the three function types. Note that the computational pipeline assigns the function type without leveraging the known biological functions. Instead, we validate these assignments against the knowledge about biological functions.

### Illustration of our computational pipeline using synthetic data

To illustrate the pipeline’s workflow, we utilized synthetic data generated by a consumer-resource model (CRM). Each niche (or specialist) function is modeled as the consumption of a unique and externally supplied resource (Fig. 2a1), whose loss would make a species unable to consume the corresponding resource (Fig. 2a2, a3). The loss of an essential function is modeled as reducing a species’ growth rate by 5% (Fig. 2a4).

**Figure 2:**
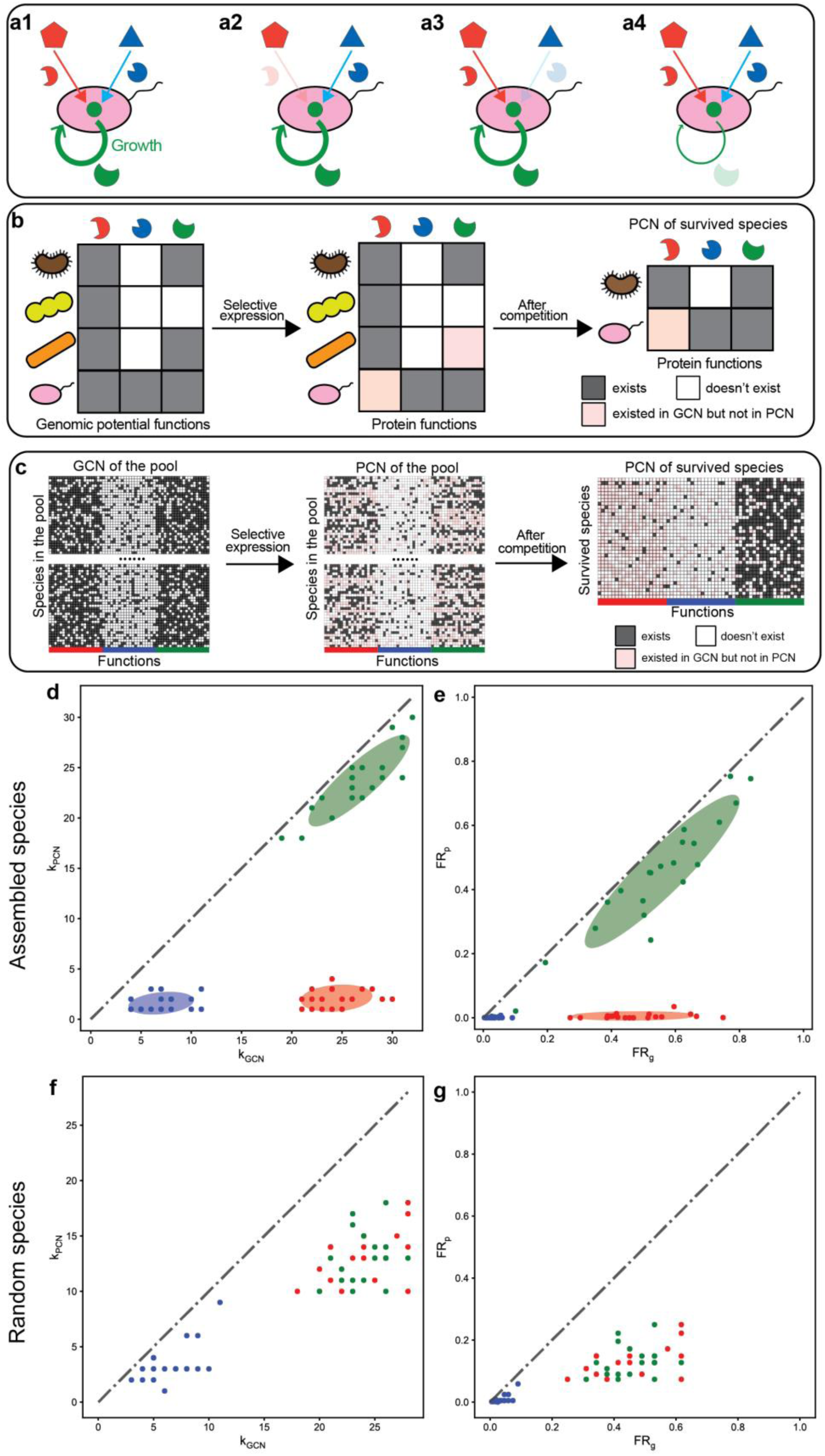
Three protein functional clusters (specialist function, essential function, and niche function) considered in the community assembly model form three distinct clusters when the network degree and functional redundancy are compared between the GCN and PCN in model-generated synthetic data. **a1-a4,** Three types of functions modeled have different ecological and metabolic roles. The niche function (red proteins) and specialist function (blue proteins) are modeled as abilities to consume externally supplied resources. The role of essential functions (green proteins) is considered as a reduction in the overall growth rate for each missing essential function. **b,** A schematic diagram of the community assembly. Species (ovals and indented ovals) with expressed gene functions selected via the sub-sampling of their genomic capacity. Then all species are co-cultured together to simulate their ecological competition. **c,** A simulation example of the community assembly, and the construction of GCN and PCN for the survived species. **d-e,** The comparison of network degree and functional redundancy respectively based on the GCN and PCN of survived species in the simulation example in panel-c. A Gaussian mixture model with 3 clusters is used to identify 3 protein functional clusters. Ellipses around clusters cover areas one standard deviation away from their means. **f-g,** The comparison of network degree and functional redundancy respectively based on the GCN and PCN of 35 species randomly selected from the 10,000 species in the initial pool. All points/functions are colored red (niche functions), green (essential functions), and blue (specialist functions) according to their types of functions in the model. k_GCN_ (or k_PCN_) is the network degree of each function in the GCN (or PCN). FR_g_ (or FR_p_) is the functional redundancy of each function on the gene level (or protein level), respectively.

For each species, each niche (specialist, or essential) function was assigned to the species’ genome with probability *p*_n_ (*p*_s_, or *p*_e_), respectively (Fig. 2b, left). We set *p*_n_ = *p*_e_ = 0.7 to ensure that we cannot distinguish niche functions from essential functions only based on their k_GCN_. We set *p*_s_ = 0.2 < *p*_n_ = *p*_e_ so that specialist functions were assigned to fewer species than niche and essential functions. Species’ actual expressed functions were determined by randomly sampling a subset of its potential functions (Fig. 2b, middle). This behavior of sub-sampling was observed when we cultured single microbial strains in different environments (Supplementary Fig. 21). We simulated community dynamics until reaching a steady state, for which we constructed the PCN of the surviving species (Fig. 2b, right; see Supplementary information for technical details). More technical details of CRM are in Supplementary information.

In our model with 10,000 species and 20 functions for each of the three function types, each species randomly sampled a subset of potential functions (Fig. 2c, left) to express (Fig. 2c, middle). We demonstrated a simulation example with 35 species surviving in the final steady state after the community assembly initialized with 10,000 species (Fig. 2c, right).

We applied the taxonomic profile, GCN, and PCN for the surviving 35 species to our computational pipeline, finding that the three modeled protein function types were correctly classified as three clusters (60 out of 60 were correct) by the Gaussian mixture model in both the comparison of network degree (Fig. 2d) and FR (Fig. 2e). We emphasize that the observed three functional clusters arise from community assembly. When we randomly picked 35 species (same as the number of surviving species) from the initial pool with equal abundances without assembly, niche functions cannot be distinguished from essential functions (Fig. 2f, g). Even when assigning the same randomly picked 35 species with the same abundances of surviving species, we still cannot differentiate these two functional types (Supplementary Fig. 2). Our findings held when varying (1) the number of species and functions or (2) model parameters *p*_n_, *p*_s_, and *p*_e_, affirming the method’s robustness in distinguishing function types (Supplementary Figs. 3, 4, and 5).

### Three protein functional clusters observed in human gut microbiomes

Next, we validated our computational pipeline on real data of human mucosal-luminal interface samples previously collected from the ascending colon of four children^14,33^. Here we focused on the genus level and annotated the identified proteins from metagenomic and metaproteomic data via the clusters of orthologous genes (COGs) database^46,47^. We chose the genus level due to widely shared peptide sequences across species (Supplementary Fig. 6). We searched metagenomic reads and metaproteomic peptides against the integrated gene catalog (IGC) database of the human gut microbiome^48^ to generate the GCN and PCN^14^, and took the intersected COGs between the two networks. Taxonomic assignment was performed using the “protein-peptide bridge” method as described previously^14^. Our analysis centers on subject HM454, identifying 1,542 intersected COGs in both the GCN and PCN and obtaining a taxonomic profile of 85 genera using MetaPhlAn2^49^. The connectance (i.e., the number of edges divided by the maximal number of possible edges) of the GCN (or PCN) is 0.220 (or 0.049), respectively (Fig. 3a, b). The GCN displayed a higher nestedness (nestedness metric NODF^50^=0.667) than the PCN (NODF=0.453). More details about data processing and NODF are in Supplementary Information.

**Figure 3:**
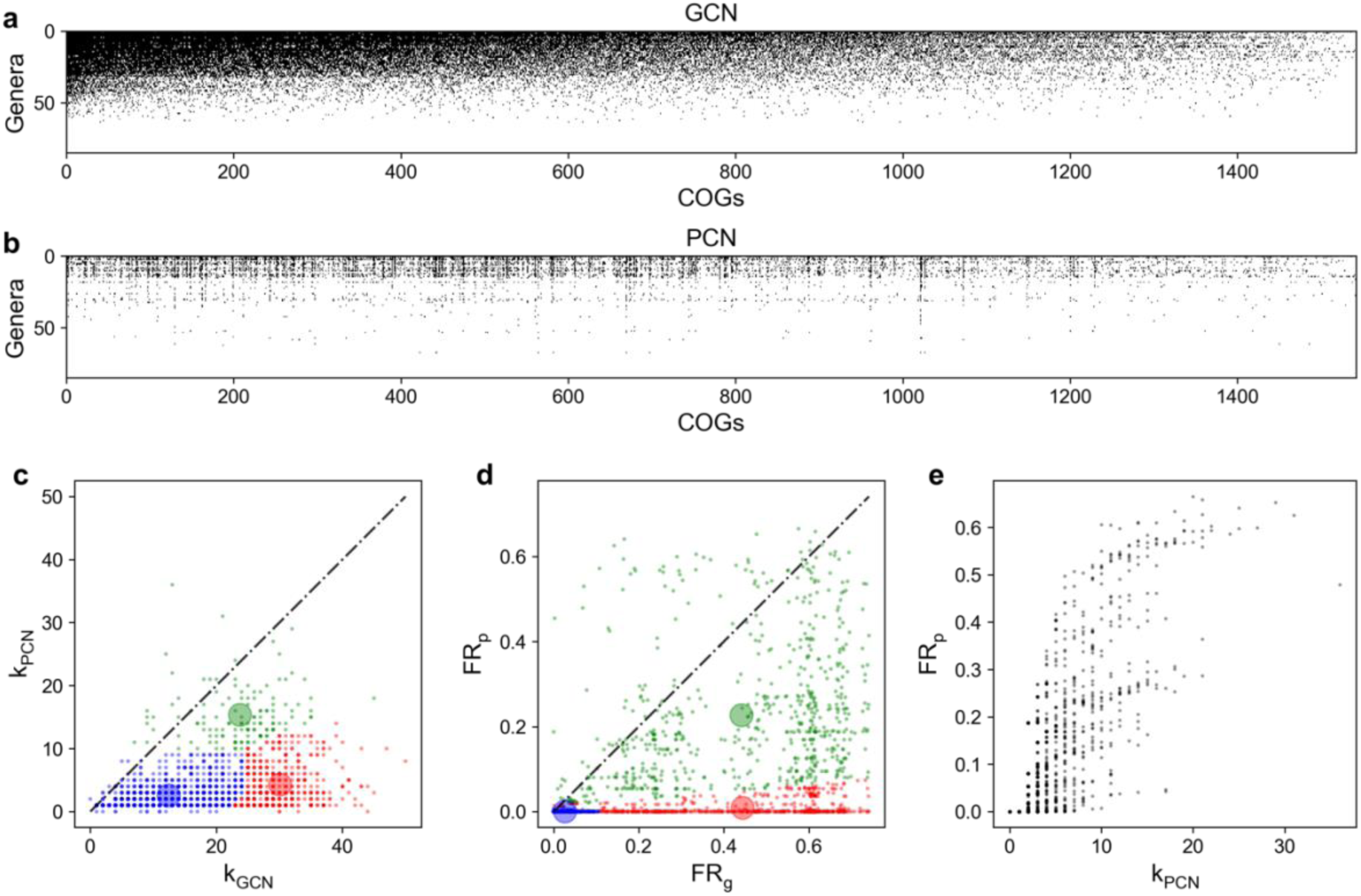
Real data of the human gut microbiome showing three clusters on the plot that compares FR_g_ with FR_p_. Metagenome and metaproteome of subject HM454 mucosal-luminal interface samples^33^ were used to construct GCN and PCN, respectively. **a,** The GCN shows if a genus owns (or doesn’t own) a COG as its genomic capacity, which is colored in black (or white). The GCN matrix is ordered to have decreasing network degrees for both genera and COGs. **b,** The PCN shows if a genus expresses (or doesn’t express) a COG as its protein function, which is colored in black (or white). The PCN matrix follows the same order as the GCN. **c,** Differences in network degree for most COGs are large. k_GCN_ is the network degree of each COG in the GCN (i.e. the number of genera owning each COG in the GCN). k_PCN_ is the network degree of each COG in the PCN (i.e. the number of genera owning each COG in the PCN). **d,** FR_g_ is larger than FR_p_ for most COGs. Three clusters with three distinct colors (blue, red, and green) are predicted by the Gaussian mixture model with 3 clusters fitted on synthetic data. The transparent large circles represent centroids of three clusters. **e,** The relationship between FR_p_ and network degree of PCN for COGs is not monotonic.

The network degree analysis revealed a general decline from GCN to PCN, with 804 out of 1,542 COGs having k_PCN_ < 0.2k_GCN_ (Fig. 3c). This decline greatly influences FR but does not fully explain why many COGs have FR_p_∼0 (744 out of 1,542 have FR_p_ < 0.01 in Fig. 3d) and k_PCN_ nonlinearly correlates with FR_p_ (Fig. 3e). For example, for L-arabinose isomerase (COG2160), its k_PCN_ (7) is fairly close to k_GCN_ (8), but its FR_p_ (0.04) is much lower than FR_g_ (0.23) since the genus *Blautia* (relative abundance = 22%) did not express L-arabinose isomerase, even if it has this capacity encoded in its genome.

Using the Gaussian mixture model fitted on simulated data, we categorized all protein families into three clusters (Figs. 3c&d). Although clusters on real data are not as distinct as on simulated data, the relative positioning of the three clusters (shaded areas in Fig. 3c, d) agrees well with our hypothesis (Fig. 1). The weaker clustering might result from a greater variation in k_GCN_ (or FR_g_) for real data (Fig. 3c, d) than that for simulated data (Fig. 2d, e).

Some COGs have FR_p_ > FR_g_ (Fig. 3c, d), contradicting the sub-sampling argument for the gene expression. FR_p_ should not exceed FR_g_ if the PCN was a proper subgraph of the GCN. This contradiction may stem from limitations in metagenomic sequencing and metaproteomic identification depths, as both metagenomics and metaproteomics require sufficient depth to detect genes or proteins, respectively. We tested how the lower detection capability of metaproteomics or metagenomics influences the FR by varying the protein or gene abundance percentile threshold (PAPT or GAPT), which denotes the percentage of most abundant proteins or genes being kept. As PAPT or GAPT decreases, FR_p_ or FR_g_ drops respectively (Supplementary Fig. 15). When GAPT decreases, we observed more proteins with FR_p_ greater than their FR_g_.

### Validating three functional clusters observed in human gut microbiomes

Our computational pipeline accurately assigns functional clusters for protein families, agreeing with their known biological functions. For example, COG0539 (ribosomal protein S1) was assigned as the essential function, which is essential for translational initiation^17–19,51,52^. Another example is the assignment of COG1116 (ABC-type nitrate/sulfonate/bicarbonate transport system)^34^ as a niche function, whose expression has been shown to be selectively enriched for a few microbial species^53^.

The pipeline’s classifications were systematically validated against well-established biological roles of specific protein families: (1) ABC-type transporters are niche proteins due to their connection with ecological metabolic niches^34–36;^ (2) Ribosomal proteins are essential proteins because they are indispensable for the microbial growth^54,55^; (3) PTS (phosphotransferase system) proteins are specialist proteins because an evolutionary study has shown that various species within the same genus even possess a different set of PTS proteins^56^. To evaluate our pipeline’s performance, we quantified its accuracy in assigning these protein families (ABC-type transporters, ribosomal proteins, or PTS proteins) against the assumed “ground-truth” functions (niche, essential, or specialist functions, respectively).

For HM454, our computational pipeline based on the FR_g_/FR_p_ plot correctly categorizes 81 of 122 COGs belonging to ABC-type transporters, ribosomal proteins, or PTS proteins. In comparison, when classifying functions based on the k_GCN_/k_GCN_ plot, 74 COGs were correctly assigned, slightly worse than that based on the FR_g_/FR_p_ plot. Specifically, 26 of 53 COGs belonging to ABC-type transporters are classified as niche functions. The fraction of ribosomal proteins classified to the cluster of essential functions is 83.0%(=44/53). For the PTS proteins, among the identified 16 COGs, 11 are classified as specialist functions.

### Alternative clustering and classification methods

Alternatively, we explored the unsupervised K-mean clustering with K=3, which captured the three representative functional clusters with their positions agreeing with our expectations (Supplementary Fig. 13). We also designed a supervised classifier based on quadratic discriminant analysis (QDA). QDA, trained on ABC-type transporters, PTS proteins, and ribosomal proteins as the niche, specialist, and essential functions, generated clusters closely resembling those from the Gaussian mixture model (Supplementary Fig. 14). For HM454, the K-mean clustering categorizes 48 of these 122 COGs that are ABC-type transporters, ribosomal proteins, or PTS proteins into clusters respectively representing niche, essential, or specialist functions (i.e., the accuracy is 39.3%). For the QDA classifier, the accuracy is 59.0%(=72/122). Thus, we select the Gaussian mixture model as the classification method because of its superior accuracy (66.4%=81/122).

### Comparing FR_g_ with FR_p_ identifies ecological niches and metabolic essentiality

We focused on analyzing ABC-type transporters^34–36^ and ribosomal proteins^17–19^. ABC-type transporters are energy-requiring transporter proteins that allow microbes to exploit specific niches like glucose uptake^34–36^. For HM454, we indeed found that k_GCN_ for all ABC-type transporters are much larger than their k_PCN_ (Fig. 4a). Similarly, we also found that their FR_g_ are much larger than their FR_p_, classifying many transporter proteins as niche functions (Fig. 4b). Some transporter proteins were classified as specialist functions (blue dots in Fig. 4b) due to the specialization on the gene level, which is carried to the protein level. Some transporter proteins were classified as essential functions (green dots in Fig. 4b). One example is the ABC-type Fe^3+^/spermidine/putrescine transporter (COG3842), as iron is essential for bacteria to function as a co-factor in iron-containing proteins^57,58^.

**Figure 4:**
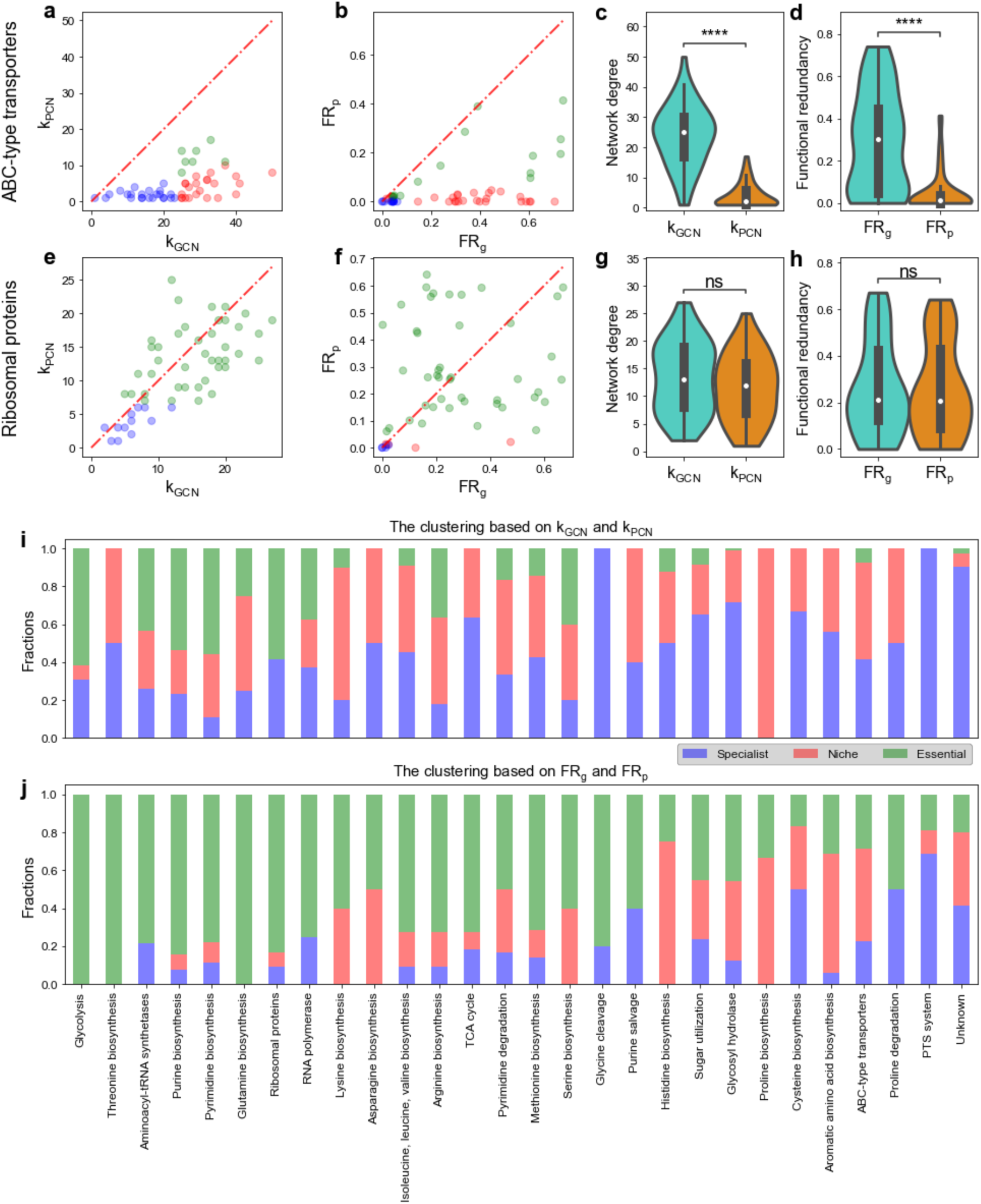
Comparison of network degree and functional redundancy between the gene and protein level for many protein families including ABC-type transporters and ribosomal proteins from the human gut microbiome. **a,** Network degrees in GCN are larger than network degrees in PCN for most ABC-type transporter COGs. k_GCN_ (or k_PCN_) is the network degree of each COG in the GCN (or PCN). **b,** FR_g_ is larger than FR_p_ for most ABC-type transporter COGs. **c-d,** The distribution of network degrees and functional redundancies (violin plots and boxplots) for ABC-type transporter COGs show a significantly huge reduction from k_GCN_ to k_PCN_ or from FR_g_ to FR_p_. **e,** Network degrees in GCN are comparable with that in PCN for most ribosomal protein COGs. **f,** FR_g_ is comparable with FR_p_ for most ribosomal protein COGs. Points in scatter plots are colored by the same colors used in Fig. 3d. **g-h,** The distribution of network degrees and functional redundancies (violin plots and boxplots) for ribosomal protein COGs show no significant reduction from k_GCN_ to k_PCN_ or from FR_g_ to FR_p_. **i,** The fraction of assigned specialist, niche, or essential functions based on comparing network degrees k_GCN_ and k_PCN_ for many protein families. **j,** The fraction of assigned specialist, niche, or essential functions based on comparing functional redundancies FR_g_ and FR_p_ for many protein families. In all boxplots, the middle white dot is the median, the lower and upper hinges correspond to the first and third quartiles, and the black line ranges from the 1.5 × IQR (where IQR is the interquartile range) below the lower hinge to 1.5 × IQR above the upper hinge. All violin plots are smoothed by a kernel density estimator and 0 is set as the lower bound. All statistical analyses were performed using the two-sided Mann-Whitney-Wilcoxon U Test with Bonferroni correction between genomic capacity (GCN) and protein functions (PCN). P values obtained from the test is divided into 5 groups: (1) *p* > 0.05 (ns), (2) 0.01 < *p* ≤ 0.05 (*), (3) 10^−3^ < *p* ≤ 0.01 (**), (4) 10^−4^ < *p* ≤ 10^−3^ (***), and (5) *p* ≤ 10^−4^ (****). Network degree comparison of ABC transporters: *p* = 7.11 × 10^−16^. Network degree comparison of ribosomal proteins: proteins: *p* = 0.10. Redundancy comparison of ABC transporters: *p* = 2.19 × 10^−11^. Redundancy comparison of ribosomal proteins: *p* = 1.00.

Ribosomal proteins, critical for protein synthesis and microbial growth^54,55^, showed little variance between k_GCN_ and k_PCN_ (the mean and the standard deviation of the relative difference 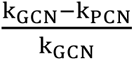 is 0.09 ± 0.41; Fig. 4e), with most correctly classified as essential (44 out 53 COGs in Fig. 4f). Notably, two ribosomal proteins (L28 and L34), which have been reported as non-essential to microbes such as *E. coli*^17,52,59^, were accurately classified as non-essential proteins (red dots in Fig. 4e). Certain specialized ribosomal proteins in microbial genomes continue to be specialized on the protein level and thus were classified as specialist functions.

Alternatively, we looked at the distribution of network degrees (Figs. 4c&g) and FR (Figs. 4d&h). For ABC-type transporters, the distribution of k_PCN_ is close to 0 (median of 2), while the median of k_GCN_ is 25. For ribosomal proteins, the distribution of k_PCN_ (median is 12) is similar to k_GCN_ (median is 14). For ABC-type transporters, the distribution of FR_p_ is close to 0 (with a median ∼ 0.01), while the median of FR_g_ is around 0.30. For ribosomal proteins, the distribution of FR_p_ (median ∼ 0.20) is similar to the distribution of FR_g_ (median ∼ 0.21).

The similar patterns are also true for the other three individuals (Supplementary Figs. 8-10). Variations in the FR_g_ /FR_p_ plot across individuals (Supplementary Fig. 8-10) are likely due to differences in gut environments, diets, and microbial composition. Note that HM503 is an outlier due to its lower diversity (36 genera versus the average of 56.7) (Supplementary Fig. 11). Similarly, the Shannon diversity index for HM503 is only 1.41, lower than other individuals (mean and standard deviation are 2.05 and 0.18). The lower diversity in HM503 leads to fewer taxa owning the same function on average, resulting in lower FR_g_ and FR_p_ values.

Extending our analysis to additional protein families, we discovered that proteins linked to glycolysis, RNA polymerase, and the TCA cycle displayed patterns similar to ribosomal proteins (Figs. 4i&j). By contrast, proteins associated with sugar utilization, glycolyl hydrolase, and aromatic amino acid biosynthesis exhibited patterns more akin to ABC-type transporters. We also analyzed the classification results for the unknown COG functions (with the category ‘S: Function Unknown’). Out of the 104 unknown COGs in our analysis, the majority are classified as specialist (43/104) or niche functions (40/104), with a smaller portion identified as essential functions (21/104).

We also confirmed our results using the KEGG Orthology (KO) annotation^60–63^, which has a lower annotation rate (78%) than COG (92%). For HM454, our computational pipeline categorizes 75 of these 126 KOs that are ABC-type transporters, ribosomal proteins, or PTS proteins into clusters respectively representing niche, essential, or specialist functions, similar to the classification accuracy based on the COG. The contrasting difference between ABC-type transporters and ribosomal proteins is well preserved (see Supplementary Fig. 7). Additionally, the distribution of FR_p_ shows a dramatic difference across KO groups (Supplementary Fig. 12). Some ecologically strongly selected KO groups such as ABC transporters have small FR_p_ (Supplementary Fig. 12). As a comparison, proteins from aminoacyl-tRNA biosynthesis^15,16^, glycolysis^20,21^, and ribosomes^17–19^ have large FR_p_ and huge variations within each group (Supplementary Fig. 12).

### Validating our method on the mouse gut microbiome

In testing our method’s feasibility in other microbial communities, we leveraged a metaproteomic dataset from mice gavaged with a synthetic microbiome comprising twenty sequenced bacteria^41^. Since this study lacks paired metagenomes, we used its 16S rRNA gene sequencing data with individual genomes to infer its metagenome. Here we focused on the strain level because peptides of different strains in this simple synthetic gut microbiome can be distinguished. We relied on the comparison between FR_g_ and FR_p_ to generate the distribution of functional clusters across many protein families for this dataset (Fig. 5n). The results mirrored those of human gut microbiomes, especially the contrasting patterns between ABC-type transporters and ribosomal proteins (Fig. 5).

**Figure 5:**
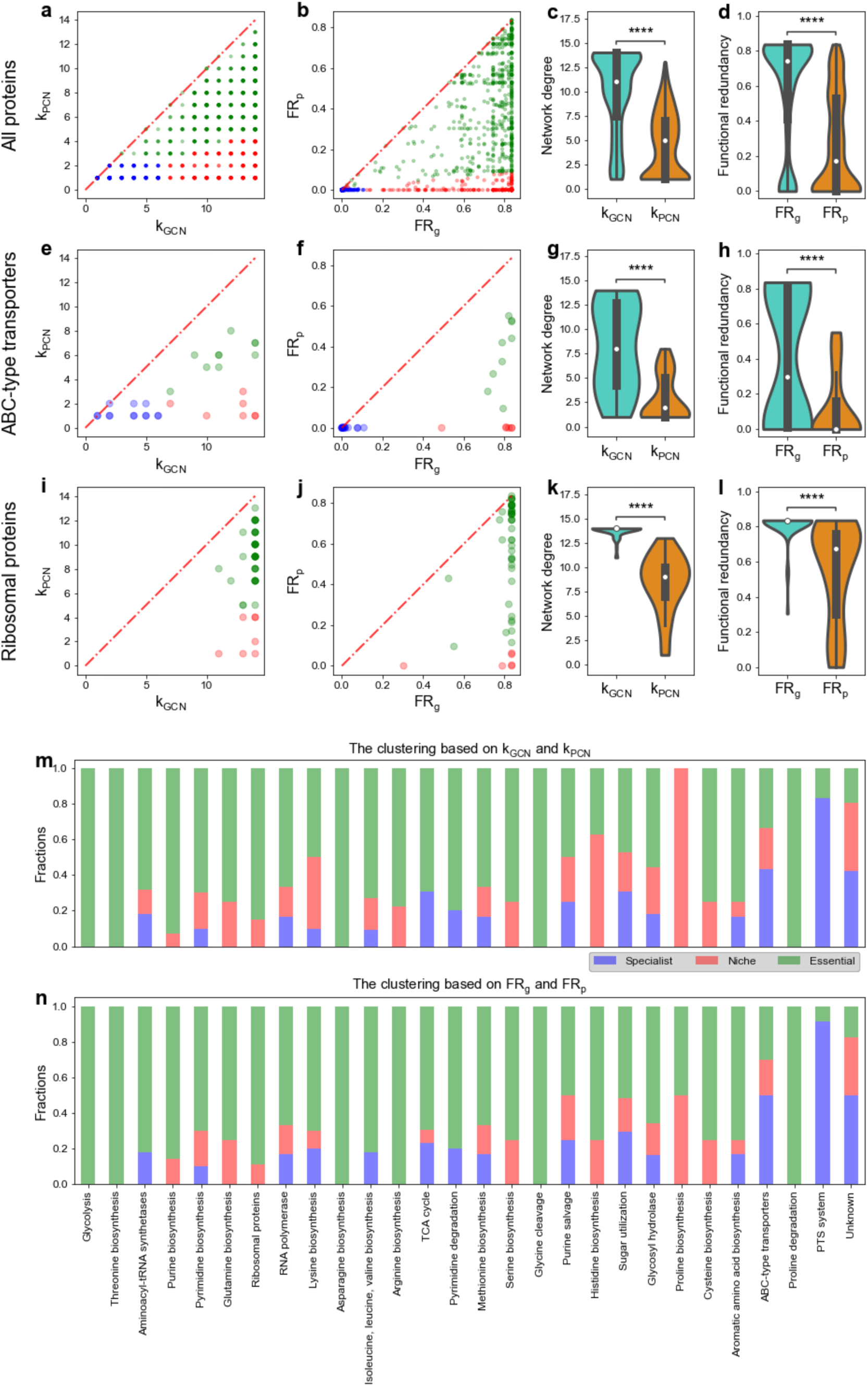
Comparison of network degree and functional redundancy between the gene and protein level for many protein families including ABC-type transporters and ribosomal proteins from the synthetic mouse gut microbial community. **a,** Comparison between network degrees of all COGs in the GCN (k_GCN_) and network degrees of all COGs in the PCN (k_PCN_). **b,** Comparison between functional redundancies of all COGs in the GCN (FR_g_) and functional redundancies of all COGs in the PCN (FR_p_). **c-d,** The distribution of network degrees and functional redundancies (violin plots and boxplots) for all COGs. **e-h,** Comparison between k_GCN_ and k_PCN_, comparison between FR_g_ and FR_p_, the distribution of network degrees, and the distribution of functional redundancies for ABC-type transporter COGs. **i-l,** Comparison between k_GCN_ and k_PCN_, comparison between FR_g_ and FR_p_, the distribution of network degrees, and the distribution of functional redundancies for ribosomal protein transporter COGs. **m,** The fraction of assigned specialist, niche, or essential functions based on comparing network degrees k_GCN_ and k_PCN_ for many protein families. **n,** The fraction of assigned specialist, niche, or essential functions based on comparing functional redundancies FR_g_ and FR_p_ for many protein families. In all boxplots, the middle white dot is the median, the lower and upper hinges correspond to the first and third quartiles, and the black line ranges from the 1.5 × IQR (where IQR is the interquartile range) below the lower hinge to 1.5 × IQR above the upper hinge. All violin plots are smoothed by a kernel density estimator and 0 is set as the lower bound. All statistical analyses were performed using the two-sided Mann-Whitney-Wilcoxon U Test with Bonferroni correction between genomic capacity (GCN) and protein functions (PCN). P values obtained from the test is divided into 5 groups: (1) *p* > 0.05 (ns), (2) 0.01 < *p* ≤ 0.05 (*), (3) 10^−3^ < *p* ≤ 0.01 (**), (4) 10^−4^ < *p* ≤ 10^−3^ (***), and (5) *p* ≤ 10^−4^ (****).

### Response of community and protein abundance to the introduction of sugars

After identifying niche functions through our computational pipeline, we explored using nutrients associated with niche functions to manipulate the community structure. In ecology, a niche is often defined as an abiotic and biotic factor that supports the survival of species^9,64–66^. Therefore, niche functions are associated with corresponding limiting resources involved in those functions. For example, COG1879 (ABC-type sugar transporter) is categorized as a niche function due to microbial competition for sugars. Here, we leveraged the *in vitro* community and studied how expression levels of ATP-type transporters respond to supplied sugars so that a microbial taxon can achieve a better living strategy.

Using the RapidAIM V2.0 approach^67^, which replicates the functional profiles of individual gut microbiomes *in vitro*^37^, we cultured three individual human gut microbiota samples and used a semi-automated metaproteomics workflow to observe how taxon-specific proteins respond to the presence of glucose, fructose, and kestose (Fig. 6a). Samples were cultured in technical triplicates, and protein abundances were quantified at 0, 1, 5, 12, and 24 hours using 11-plex tandem mass tag (TMT11plex)^39^ for a total of 189 samples. We analyzed the Bray–Curtis dissimilarity of metaproteomes over time, finding that more complex sugars induce more pronounced alterations in protein profiles (Supplementary Fig. 16).

**Figure 6:**
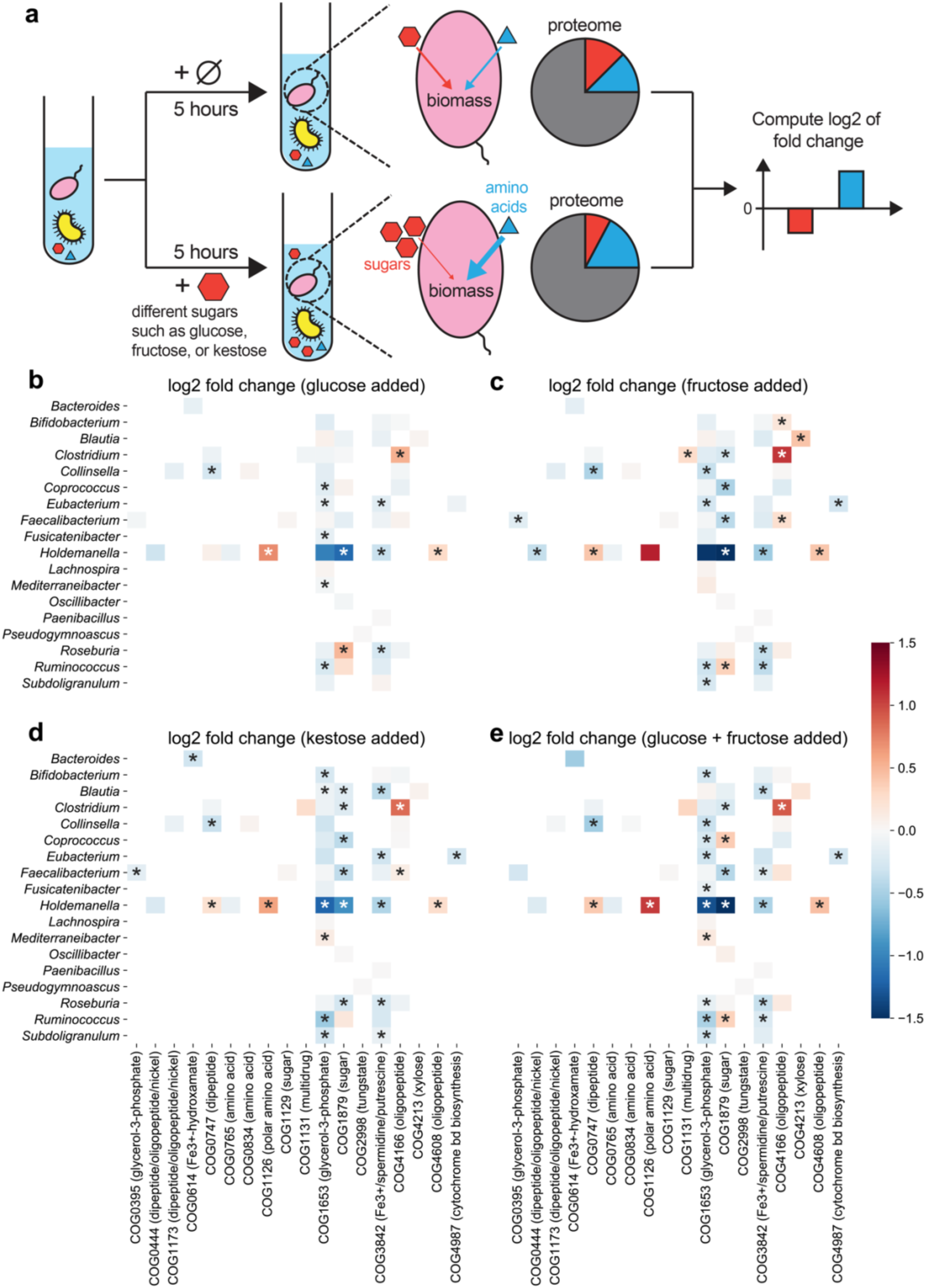
Microbes modify their expression for ABC-type transporters to adapt to added sugars. All heatmaps share the same color bar on the right. **a,** Schematic of *in-vitro* cultures of a collected human gut microbiome. In the treatment group, one sugar is added to the community. Metaproteomic measurements 5 hours later were used to compare the intensity of each taxon-specific protein using the log2 fold change of each protein’s fraction (i.e. normalized intensity over each genus) from the treatment group divided by that from the control group. Log2 fold changes of ABC-type transporters were computed 5 hours after (**b**) glucose, (**c**) fructose, (**d**) kestose, or (**e**) glucose and fructose is added. The transported metabolites for each COG are added to the brackets.

To reflect the effect of introduced sugars on protein abundances, we used log2 of fold change in normalized protein abundances/intensities (see Supplemental Methods for details) between the treatment and control group (Fig. 6). We hypothesized that excessive sugars remove the growth limitation on carbon resources, prompting microbes to upregulate other transporters for uptaking more other scarce resources (e.g., nitrogen or amino acids) for better growth (Fig. 6a). We analyzed log2 fold changes of ABC-type transporters 5 hours later after sugar introduction (Figs. 6b-d), with most COGs close to zero. Among seven significantly influenced COGs, COG1126 (ABC-type polar amino acid transport system) is the only one that is revealed to be a niche function. Focusing on COG1126, we found that it is specialized by the genus *Holdemanella*. *Holdemanella* benefits from upregulating COG1126, as the proportion of *Holdemanella* proteins significantly increases from 13.5%(± 0.06%) for the control to 15.8 %(± 0.08%) with the added glucose (p-value = 0.04, Mann-Whitney U test applied).

We observed that adding fructose, glucose and fructose, or kestose alters ABC-type transporters’ expression similarly to glucose alone (Fig. 6). The correlation in log2 fold changes of ABC-type transporters between different added sugars is significant (Supplementary Fig. 18). Notably, complex sugars trigger more significant fold changes in microbial protein expression. This pattern persists for metaproteomic measurements 12 and 24 hours later, while the fold changes 1 hour later are less significant (Supplemental Fig. 19; p-value < 0.01 for four sugar-adding scenarios, Mann-Whitney U test applied). Additionally, the overwhelmingly positive log2 fold changes of ribosomal proteins (Supplementary Fig. 20; p-value < 10^-4^ for four sugar-adding scenarios, one-sample Wilcoxon test applied) probably implies faster microbial growth when simple sugars are supplied^68,69^.

## Discussion

We developed a computational pipeline to classify protein families as specialist, niche, and essential functions by comparing FR_g_ with FR_p_. This approach supplements traditional methods that test metabolic essentiality by gene knockout^17–19^ and identify limiting resources by measuring biomass changes upon resource supplies^70–73^. We first illustrated this method on synthetic data and then validated it using real datasets of human and mouse gut microbiomes. We acknowledge a limitation in our validation process—the reliance on limited available literature. Hence, our classification should be seen as a preliminary framework that is open to refinement as the investigation of protein families’ functions improves.

Our findings bridge the gap between the ecological niche theory, which posits that each resource (or niche) can only be occupied by one species for steady-state conditions^66,74,75^, and the FR revealed by shared functions among microbial genomes^11,12^. We solved this dilemma by showing niche proteins usually have very small FR_p_ and large FR_g_. Additionally, our ecological framework combines genomic capacity and protein functions together by introducing species with sub-sampled functions. The model framework accounts for selective expression due to different environmental conditions^76–78^ or evolved strains with distinct metabolic niches^79,26,27^, reconciling phenotype-focused ecological models with genetic data.

The observed case of FR_p_ > FR_g_ for some COGs could stem from using MetaProIQ, a general microbiome catalog, for metaproteome analysis. Although MetaProIQ facilitates the identification of proteins from various gut microbes, the general search against it may cause anomalies. In contrast, directly searching metaproteomic data against gene calls from the paired metagenome may deliver more accurate identifications. However, this strategy suffers when the metagenomic sequencing is incomplete, leading to undetected proteins due to missing genes. In our human gut microbiome datasets, limited sequencing depth might cause such issues. For the synthetic mouse gut communities with complete genomes for each microbial strain, we matched metaproteome to microbial genomes and did not encounter any case of FR_p_ > FR_g_, highlighting the effectiveness of this approach in contexts with complete genomic data.

In this work, we only validated our pipeline on gut-related biomes due to the limited accessibility of paired metagenome-metaproteome in other environments. In nutrient-poor environments, where ribosomal proteins are less expressed^80,81^, essential proteins like ribosomes may be harder to detect, causing potential detection biases. This aspect warrants further investigation to ensure the robustness and applicability of our method in diverse ecological settings. Other technical limitations can also impact clustering accuracy. Smaller ribosomal proteins L28 and L34 could be detected less frequently in metaproteomics, and post-translational modifications may also result in missed cleavage and identification of peptides. Advances such as metaproteomics-assembled proteomes (MAPs)^82^ may improve taxon-specific functional annotations and the accuracy of our clustering outcomes.

## Data and code availability

All code for simulations used in this manuscript can be found at https://github.com/wt1005203/ecological_niches.

## Acknowledgements

We thank Dr. Janice Mayne for the help with providing biobanking samples, Dr. Zhibin Ning for the help with running mass spectrometry and Dr. Joeselle Serrana for the help with processing the metagenomic data. D.F. acknowledges grants from Natural Sciences and Engineering Research Council of Canada (NSERC), and the Government of Canada through Genome Canada and the Ontario Genomics Institute (OGI-114 & OGI-149). D.F. acknowledges a Distinguished Research Chair from the University of Ottawa. Y.-Y.L. is supported by grants R01AI141529, R01HD093761, RF1AG067744, UH3OD023268, U19AI095219 and U01HL089856 from the National Institutes of Health, USA; a pilot grant from the Biology of Trauma Initiative of Broad Institute, USA; and the Office of the Assistant Secretary of Defense for Health Affairs, through the Traumatic Brain Injury and Psychological Health Research Program (Focused Program Award) under award no. (W81XWH-22-S-TBIPH2), endorsed by the Department of Defense, USA.

## Author contributions

Y.-Y.L. and D.F. supervised the study. T.W. and Y.-Y.L. conceived the project. All authors designed the research. L.L. prepared and curated the empirical data as well as performed all wet-lab experiments. T.W. analyzed all data and developed the ecological model. T.W. wrote the initial manuscript. All authors edited and approved the manuscript.

## IRB determination/approval statement

The sample collection of human gut microbiomes was approved by the Research Ethics Board of the Children’s Hospital of Eastern Ontario (CHEO), Ottawa, ON, Canada. The written informed consent forms were obtained from their parents.

