## Supplemental Information for "Pairing Metagenomics and Metaproteomics to Characterize Ecological Niches and Metabolic Essentiality of gut microbiomes"

### Supplementary Information

#### Supplemental Methods

***In-vitro* culture of single gut bacterial strains with added sugars.** Five gut commensal bacterial strains, *Bacteroides vulgatus* ATCC 8482, *Bacteroides ovatus* ATCC 8483, *Bacteroides uniformis* ATCC 8492, *Blautia hydrogenotrophica* DSM 10507, *Escherichia coli* DSM 101114 were cultured with or without added sugars (glucose, sucrose, and kestose). The base culture medium without sugar added were modified based on the Yeast Casitone Fatty Acids (YCFA) broth, containing 10.0 g/L casitone, 2.5 g/L yeast extract, 45 mg/L $MgSO_4 \cdot 7H_2O$, 90 mg/L $CaCl_2 \cdot 2H_2O$, 450 mg/L $K_2HPO_4$, 450 mg/L $KH_2PO_4$, 900 mg/L NaCl, 1.0 mg/L resazurin, 4.0 g/L $NaHCO_3$, 1.0 g/L L-Cysteine-HCl, 10 mg/L Hemin, 1.90 mL/L acetic acid, 0.7 mL/L propionic acid, 90 μL/L iso-butyric acid, 100 μL/L n-valeric acid, 100 μL/L iso-valeric acid, 0.02 mg/L biotin, 0.02 mg/L folic acid, 0.05 mg/L thiamine-HCl, 0.05 mg/L riboflavin, 0.001 mg/L vitamin $B_{12}$, 0.05 mg/L aminobenzoic acid. The pH was adjusted to between 6.7-6.8, and autoclaved media were pre-reduced in an anaerobic chamber overnight. 5 g/L of different sugars (glucose, sucrose, and kestose) were added to the base medium as treatment groups. Master tubes of single bacterial strains were first cultured on Tryptic Soy Agar (TSA) containing 5% sheep blood using the streak plate method. A single colony was picked from each agar plate and inoculated into the base culture medium to culture for 24 hours, before inoculating 100 μL of each culture into 10 mL of four different media: base medium without sugar added, with glucose added, with sucrose added and with kestose added. After culturing for 24 hours, optical density at 600 nm was tested in technical triplicates for each sample. Cultured microbial cells were purified by washing with phosphate buffered saline (PBS) buffer three times, and the resulting microbial pellets were stored at -80 ºC for proteomics analysis.

***In-vitro* human gut microbiota culture with added sugars.** Three healthy individual microbiota samples were collected and biobanked using our live microbiota biobanking protocol[83]. The study was approved by the Ottawa Health Science Network Research Ethics Board at the Ottawa Hospital, Ottawa, Canada (# 20160585–01 H). The frozen microbiome samples were thawed at 37 ºC and cultured in our optimized culture medium[37] with or without the presence of different sugars (10 mM glucose, 20 mM fructose, 10 mM glucose + 20 mM fructose, or 10 mM kestose). Samples were cultured in technical triplicates, and were taken at 0 hr, 1hr, 5 hr, 12 hr, and 24 hr of culturing for optical density and metaproteomic analyses.

After culturing, 96-well deep well plates were first centrifuged at 3,000 g for 45 min under 4 °C. Then the pellets were washed in 4 °C phosphate buffered salin (PBS) buffer and centrifuged at 3,000 g for 45 min again, before pelleting and removing culture debris three times using 300 g, 4 °C , 5 min centrifugation. Microbial suspensions were then centrifuged at 3,000 g, 4 °C for another 45 min. The purified cell pellets were stored at -80 °C before protein extraction.

**Protein extraction, digestion and LC-MS/MS analysis.** For single strain samples, proteins were extracted with 4% SDS 8M urea buffer in 100 mM Tris-HCl buffer and precipitated overnight at -20 ºC, before being purified by washing with ice-cold acetone three times. Quantified proteins were then reduced and alkylated before being digested using trypsin (50:1 protein-to-trypsin ratio) for 24 hours at 37 ºC and were desalted using reverse phase beads[38]. Proteomic samples were analyzed using an Orbitrap Exploris 480 mass spectrometer (ThermoFisher Scientific Inc.) coupled with an UltiMate 3000 RSLCnano liquid chromatography system following a 1-hour gradient of 5 to 35% (v/v) acetonitrile (v/v) at the flow rate of 300 L/min. MS full scan was performed from 350 - 1400 m/z with a resolution of 60,000, followed by an MS/MS scan of 12 most intense ions, a dynamic exclusion repeat count of one, exclusion duration of 30 s, and resolution of 15,000. Metaproteomics samples of the cultured individual microbiomes were prepared using a semi-automated approach. Briefly, samples were lysed in a buffer containing 8 M urea, 4% SDS in 100 mM Tris-HCl (pH = 8.0) to extract microbial total proteins. The proteins were purified by a double-precipitation procedure in 50%:50%:0.1% (v/v/v) acetone: ethanol: acetic acid solution. Protein digestion and desalting steps were performed using an automated liquid handler (Hamilton Nimbus-96). Briefly, 100 μg proteins were dissolved in 100 μL 6 M urea in 100 mM Tris-HCl (pH 8) buffer, before being reduced by 10 μL 0.1 M dithiothreitol (DTT) solution under 56 °C for 30 minutes and alkylated by 10 μL 0.2 M iodoacetamide (IAA) solution in dark, 25 °C for 40 minutes. Samples were each added 1000 μL 100 mM Tris·HCl buffer containing 2 μg/mL trypsin (trypsin:proteins = 1:50) for a 24-hour digestion under 37 °C, before being desalted using an automated pipeline based on reverse-phase (RP) desalting columns. 11-plex tandem mass tag (TMT11plex) was used for metaproteomic quantification for a total of 189 samples. An even mixture of all samples was used as the reference channel in each 11-plex. Samples were scrambled before labeling with TMT11plex, so that each labeled sample contains samples from different individuals, different time points and different treatments to avoid any bias that may be induced between analyses. TMT-labelled samples were analyzed using an Orbitrap Exploris 480 mass spectrometer (ThermoFisher Scientific Inc.) coupled with an UltiMate 3000 RSLCnano liquid chromatography system following a 2-hour gradient of 5% to 35% solvent B (80% acetone nitrile, 0.1% formic acid, v/v).

**Datasets.** Metagenomics data corresponding to the ultra-deep metaproteomic analysis of the four individual microbiomes were obtained from the previous MetaPro-IQ study[14,33] (accessible from the NCBI sequence read archive (SRA) under the accession of SRP068619, including "Metagenome of MLI sample in individual HM454", "Metagenome of MLI sample in individual HM455", "Metagenome of MLI sample in individual HM466", "Metagenome of MLI sample in individual HM503") and the same samples were reanalyzed by an ultra-deep metaproteomics approach[14] (accessible through the ProteomeXchange Consortium (http://www.proteomexchange.org) via the PRIDE partner repository[40] with the dataset identifier PXD027297). Proteomics dataset of the cultured singles strain samples has been deposited to ProteomeXchange Consortium via the PRIDE partner repository with the dataset identifier PXD037923. Metaproteomic dataset of the RapidAIM-cultured microbiome samples has been deposited to ProteomeXchange Consortium via the PRIDE partner repository with the dataset identifier PXD037925. The metaproteomic dataset of the mouse gut microbiome comprising twenty gut microbes is derived from a previous study[41] that was deposited to ProteomeXchange Consortium with the dataset identifier PXD009535 and to MassIVE with the dataset identifier MSV000082287.

**Database search and data processing.** Proteomics database searches were performed by combining FASTA databases of the individual strains downloaded from NCBI. The databases were combined for performing database search using MaxQuant[42] 1.6.17.0, with the label-free quantification option turned off. Metaproteomic database searches of cultured microbiome samples were performed using MetaLab V2.2[43], MaxQuant option was used to search the TMT dataset against the IGC database of the human gut microbiome. The resulting data table was normalized using R package MSstatsTMT , and missing values were imputed using R package DreamAI[45]. The "fraction" of each taxon-specific protein is computed by dividing the protein intensity by the sum of intensities of all proteins assigned to the same taxon. The log2 fold change of each protein is obtained by taking log2 of the ratio between its fraction in the treatment group (with added sugars) and its fraction in the control group (without added sugars).

**Generation of GCN and PCN.** For the ultra-deep metaproteomic dataset, the genus-COG version of GCN and PCN tables were directly obtained from the previous work[14]. In addition, here we generated a genus-KEGG version of GCN and PCN for each individual microbiome using a similar method. Briefly, for the genus-KEGG GCN, by searching raw metagenomic reads against an integrated gene catalog (IGC) database of the human gut microbiome[48], we obtained a list of proteins quantified by read counts. FASTA sequences of these proteins were searched against the KEGG database using GhostKOALA[84]. Taxonomic origination of the proteins was obtained by searching against an in-house database generated with the NCBI non-redundant (nr) database (downloaded 2/3/2016). To generate genus-KEGG PCN, the taxonomic table of the metaproteomics dataset was directly obtained from MetaLab, and KEGG annotation was also performed by querying protein FASTA sequences with GhostKOALA. Protein group intensity was used as the quantification information in PCNs. For the proteomic dataset of single strains, the whole proteomic FASTA database was submitted to EggNOG mapper (http://eggnog-mapper.embl.de/, submitted Oct-30-2021, ran emapper.py 2.1.6) to obtain functional annotations. To generate GCN, protein coding sequence (CDS) files were downloaded from NCBI, and the count of each protein id in the CDS files was considered as the copy number of each gene in the GCN. For PCN generation, intensities of identified proteins matched to each strain were used. Note that protein ids in the CDS file were 100% matched with those in the proteomic FASTA database in each strain. For the metaproteomics dataset of the cultured microbiome samples, functional information for the generation of PCN was obtained from the resulting functional table automatically generated by the MetaLab software. Taxonomic assignment was performed using the "protein-peptide bridge" method as described previously[14]. The PCNs for this dataset were then generated based on intensities of COG-genus pairs.

#### The community assembly model.

There is a longstanding gap between the ecological model which considers the protein functions of organisms and the data analysis of genomic data to give ecological insights. Ever since Robert MacArthur proposed a community model in 1970 to consider how different consumers compete exclusively for renewing resources[85], many extensions of this model were proposed to include more complex ecological factors such as cross-feeding interactions[86–89] and multiple essential nutrients[90]. Almost all of them focus on the phenotype of microbes because only functions of expressed proteins are relevant for the consumption and production of nutrients in the ecosystem. Due to the lack of metaproteomic data, many computational approaches attempting to generate ecological implications rely on the over-complete inferred protein capacity derived from genomes[7,9–12]. To reconcile this gap, we built an ecological framework with the genomic capacity and protein functions together by introducing species with sub-sampled functions. The model framework is useful for explaining the difference between genomic capacity and protein functions. The selective expression can be considered as the same microbe with different expressions under different environments[76–78] or evolved strains from the same species that have distinct metabolic niches observed in evolutionary experiments of microbes[79,26,27]. The synthetic data can be generated in four steps:

*Step 1: Assignment of species' genomic capacity.* Three types of protein functions are modeled: niche function, specialist function, and essential function. Both specialist function and niche function are considered as the capacity to consume a unique and externally supplied resource. The probability of a random consumer being assigned the ability to have a niche function is 0.7. To make fewer species own specialist functions in their genomes, the probability of a random consumer being assigned the ability to have a specialist function is 0.2, much lower than the probability of owning a niche function. The maximal consumption rate of a resource by one species represents the consumption rate that the species would have if it allocates the entire proteome (100%) to the consumption of the resource. If many resources are consumed, the total proteome has to be divided into several parts and the consumption rates would be a fraction of the corresponding maximal consumption rates. The essential function is not modeled as the consumption of alternative resources due to its metabolic essentiality. Instead, the essential function is modeled as multiplying the growth rate by a factor of 0.95 for each missing essential function.

*Step 2: Assignment of species' protein functions based on their genomic capacity.* Each species sub-samples its genomic potential functions with a sub-sampling probability $p$ (which is a random number uniformly distributed between 0 and 1) to obtain its protein functions (i.e. which resource it can truly consume). As a result, all protein functions of species form the basis for PCN. The true consumption rate of one species on a resource is its maximal consumption rate on the resource divided by the number of resources that can be utilized by the species. This process can be thought of as the proteome allocation to consume several resources simultaneously[68,69]. This assumption imposes a trade-off between a generalist and a specialist species: a generalist species utilizes more resources but has lower consumption rates for all resources, while a specialist species consumes fewer resources but has higher consumption rates for consumed resources.

*Step 3: Community assembly.* We assumed a chemostat environment, similar to the setting considered by many Consumer-Resource models[86,88]. The dilution rate $D$ is considered as 0.1 per hour. A fixed number of resources is considered and the pool concentrations (or supply rates) for all resources are assumed to be the same for simplicity. For each species, the growth rate is treated as the sum of consumption rates for different resources divided by the yield. For simplicity, all yields are assumed to be equal ($Y$ = 1). Overall, the dynamics for the concentrations of resource $i$ (denoted as $C_i$) and the abundance of the species $\alpha$ (written as $B_\alpha$):

$$\frac{dC_i}{dt} = h_i - DC_i - \frac{\sum_\beta a_{\beta i}\gamma^{N_m} B_\beta C_i}{Y}, \quad (3)$$

$$\frac{dB_\alpha}{dt} = -DB_\alpha + \sum_j a_{\alpha j}\, \gamma^{N_m} B_\alpha C_j, \quad (4)$$

where $a_{\alpha i}$ is the consumption rate of species $\alpha$ on resource $i$, $h_i$ is the supply rate of resource $i$, $Y$ is the same yield assumed for all resources, $\gamma(= 0.95)$ is the diminishing rate for the overall consumption rate that is multiplied for each missing essential function, and $N_m$ is the number of missing essential functions. The consumption rate of one species of a resource is randomly drawn from the uniform distribution between 0 and 1. Eventually, for each species, its true consumption rates are its randomly drawn consumption rates divided by the number of resources the species can use to constrain the total proteome budget[68,69]. The incidence matrix of the consumption abilities establishes part of PCN for niche functions and specialist functions of the species. The entire PCN is completed by including the presence/absence information of all essential functions.

*Step 4: Generate GCN and PCN for survived species.* When we simulated the above community assembly process to reach a steady-state in the chemostat environment, survived species can be found as species existing with non-negative abundances at the end of the simulation. For survived species, we can reconstruct the GCN and PCN for them. Within equipped GCN and PCN, we would be able to compute $\mathrm{FR_g}$, $\mathrm{FR_p}$, and network degrees ($k_{GCN}$ and $k_{PCN}$).

#### More details about validating our computational pipeline using a consumer-resource model

Previously developed consumer-resource models (CRMs) only focus on the physiologies of microbes (i.e. phenotypes)[91–93]. Simply put, those models ignored genomic capacity or potential functions, but only considered expressed functions (e.g., how species consume different resources). There was no attempt to build a consumer-resource model of microbial communities that integrates both potential and expressed functions. As a first step toward this direction, we constructed such a model.

We assumed three types of protein functions: niche functions (colored red), specialist functions (colored blue), and essential functions (colored green) in a functional pool. For simplicity, each of the niche (or specialist) functions is modeled as the consumption of a unique and externally supplied resource (Fig. 2a1). To model the difference between niche and specialist functions, we assume they are associated with different numbers of species (i.e., "consumers" in the consumer-resource modeling framework). The former should be associated with much more species than the latter. The loss of a niche or specialist function would make a species unable to consume the corresponding externally supplied resource (Fig. 2a2, a3). The loss of an essential function is simply modeled as the reduction of a species' growth rate (Fig. 2a4). Mathematically, we multiply the intrinsic growth rate of a species by a diminishing factor $\gamma = 0.95$ for each missing essential function.

The key issue in this genome-aware consumer-resource modelling framework is to decide how microbes select a subset of their potential functions to express. To tackle this issue, we first assigned potential functions to each species (Fig. 2b, left). In particular, for each species, each niche (specialist, or essential) function was assigned to the species' genome with probability $p_{\rm n}$ ($p_{\rm s}$, or $p_{\rm e}$), respectively. In our simulations, we set $p_{\rm n} = p_{\rm e} = 0.7$ to ensure that we cannot distinguish niche functions from essential functions only based on GCN and thus would like to see if they show different patterns after the community assembly. We set $p_{\rm s} = 0.2 < p_{\rm n} = p_{\rm e}$ so that specialist functions were assigned to fewer species than niche and essential functions. Then for each species, we determined its truly expressed functions by randomly sub-sampling a subset of its potential functions (Fig. 2b, middle). This behavior of sub-sampling of genomic abilities as true expressions was observed when we cultured single microbial strains in different environments (Supplementary Fig. 21). For function type-$\alpha$ ($\alpha = 1,2,3$), this was achieved by expressing each potential function with a species-specific and function-type-specific probability $p_{i,\alpha}$ randomly drawn from a uniform distribution $\mathcal{U}(0,1)$. Since different species have different sub-sampling probabilities, some species will tend to be generalists (or specialists). Similar to all consumer-resource models[91–93], we assume a fixed expression pattern for each species and all resources being supplied so that we don't have to consider the complexity of adaptive expression (such as different expression patterns when different resources are supplied). In the end, we assembled all species in the same community and ran consumer-resource dynamics until the system reached a steady state, for which we constructed the PCN of the survived species (Fig. 2b, right).

We assumed the species pool consists of $N = 10{,}000$ species, and the function pool consists of 20 functions for each of the three function types. We introduced $10{,}000$ species to ensure the number of initial species in the assembly simulation is much larger than the number of functions so that we can assemble a high-diversity community in the end. Starting from the GCN of the initial species pool (Fig. 2c, left), for each species, we randomly sub-sampled a subset of potential functions to express (middle panel, Fig. 2c). For each species, its true consumption rates are its maximal consumption rates divided by the number of resources the species can use (see Methods) to prevent the selection of generalist species that consume all resources without a penalty[68,69]. Due to the competitive exclusion principle[94], the maximal number of species surviving in the final steady state is 40, because there are 40 unique externally supplied resources ("nutrients") in our model.

We demonstrated a simulation example with 35 species surviving in the final steady state (Fig. 2c, right). For this assembled steady-state microbial community, we found that the three modeled protein functions types were correctly revealed as three clusters by the Gaussian mixture model in both the comparison of network degree (Fig. 2d) and FR (Fig. 2e). In particular, for niche functions (red cluster in Fig. 2d, e), their mean degree in PCN (2.1) is much lower than that in GCN (24.45), and their mean $\mathrm{FR_p}$ (0.005) is also much lower than their mean $\mathrm{FR_g}$ (0.48). For essential functions (green cluster in Fig. 2d, e), their mean degree in PCN (23.7) is close to that in GCN (26.7), and their mean $\mathrm{FR_p}$ (0.47) is also similar to their mean $\mathrm{FR_g}$ (0.57). For specialist functions (blue cluster in Fig. 2d, e), both their $\mathrm{k_{GCN}}$ and $\mathrm{k_{PCN}}$ (or $\mathrm{FR_g}$ and $\mathrm{FR_p}$) are low.

The three functional clusters revealed by the classification of network degrees and functional redundancies for all modeled protein functions exactly match the three function types in our model. Moreover, the relative positioning of the three functional clusters based on our simulation data agrees well with our hypothesis (Fig. 1). This clearly validates our hypothesis that niche-occupying proteins have a larger difference in FR and network degree than metabolically essential proteins.

We emphasize that the three functional clusters observed in the $\mathrm{k_{GCN}}$ vs. $\mathrm{k_{PCN}}$ (or the $\mathrm{FR_g}$ vs. $\mathrm{FR_p}$) plot is highly nontrivial. It is a result of the community assembly. To demonstrate the importance of community assembly, we randomly picked 35 species (same as the number of survived species) from the initial pool with equal abundances (i.e., the relative abundance is 1/35 for each species) without natural selection and found that it is impossible to distinguish niche functions from essential functions (Fig. 2f, g). Interestingly, for essential functions, we noticed that those species survived after the community assembly tend to have much larger $\mathrm{FR_p}$ (with mean of 0.478) than randomly selected species (with mean of 0.132). By contrast, for niche functions, survived species tend to have a smaller $\mathrm{FR_p}$ (with mean of 0.005) than randomly selected species (with mean of 0.133). Similarly, we also computed FR for the same randomly picked 35 species that share the abundances as survived species in the simulation. Again, we cannot differentiate niche functions from essential functions (Supplementary Fig. 2).

We also simulated another community with 100 niche functions, 100 specialist functions, and 100 essential functions. The species pool still consists of $N = 10{,}000$ species. The simulated results are similar to that for the community with fewer functions (Supplementary Fig. 3; Fig. 2). In addition, we tested the robustness of model parameters by varying $p_\mathrm{n}$, $p_\mathrm{s}$, and $p_\mathrm{e}$, finding that patterns of comparison of $k_{GCN}$ and $k_{PCN}$ (Supplementary Fig. 4) and comparison of $FR_g$ and $FR_p$ (Supplementary Fig. 5) are highly reproducible.

We noticed that the assumption of the trade-off between generalists and specialists (represented by assuming that the total proteome is relatively constant) is very important. In our model, this assumption is achieved by considering true consumption rates in PCN as maximal consumption rates in GCN divided by the number of resources. The importance of this trade-off lies in the fact that it forces the niche partitioning among species. In the absence of this assumption, there is no pattern of redundancy difference since generalists can always out-compete specialists. This trade-off makes sense because typically the total proteome budgets for microbes have been observed to be relatively fixed[68,69].

**Normalized gene-level functional redundancy ($\mathrm{nFR_g}$) and normalized protein-level functional redundancy ($\mathrm{nFR_p}$).** Across multiple samples, it is pointless to compare the $\mathrm{FR_g}$ or $\mathrm{FR_p}$ directly because of the difference in microbial taxonomic diversities. In fact, it has been shown in the past that the normalized functional redundancy, which is the functional redundancy divided by the taxonomic diversity, can be compared across samples[12]. In our study, the definition for $\mathrm{nFR_g}$ is

$$\mathrm{nFR_g} = \frac{\sum_{i=1}^{N}\sum_{j\neq i}^{N}\left(1-d_{ij}^{\mathrm{GCN}}\right)p_i p_j}{\sum_{i=1}^{N}\sum_{j\neq i}^{N}p_i p_j}, \tag{1}$$

and the definition for $\mathrm{nFR_p}$ is

$$\mathrm{nFR_p} = \frac{\sum_{i=1}^{N}\sum_{j\neq i}^{N}\left(1-d_{ij}^{\mathrm{PCN}}\right)p_i p_j}{\sum_{i=1}^{N}\sum_{j\neq i}^{N}p_i p_j}. \tag{2}$$

**Calculation of nestedness.** To reveal the nested structure of an incidence matrix, we first need to use the Nestedness Temperature Calculator (NTC)[95] to organize the matrix. Then we adopted the Nestedness based on Overlap and Decreasing Fill (NODF) measure previously defined[50]. The measure can only be computed for binary incidence matrices. As with any perfectly nested matrix, two properties must be present: (1) decreasing fill, which means that the columns below and to the right should have fewer entries than the columns above and to the left; and (2) paired overlap, which implies that when an entry appears in the columns below and to the right, it should also appear in the columns above and to the left. The NODF measure is calculated by averaging these two properties across all pairs of an upper and lower row and a left and right column. For the comparison of each pair, if decreasing fill is not satisfied, the pair will contribute 0 to the total nestedness. Otherwise, the pair's contribution is the percentage overlap in non-zero entries between the two rows or two columns.

### Supplementary Figures

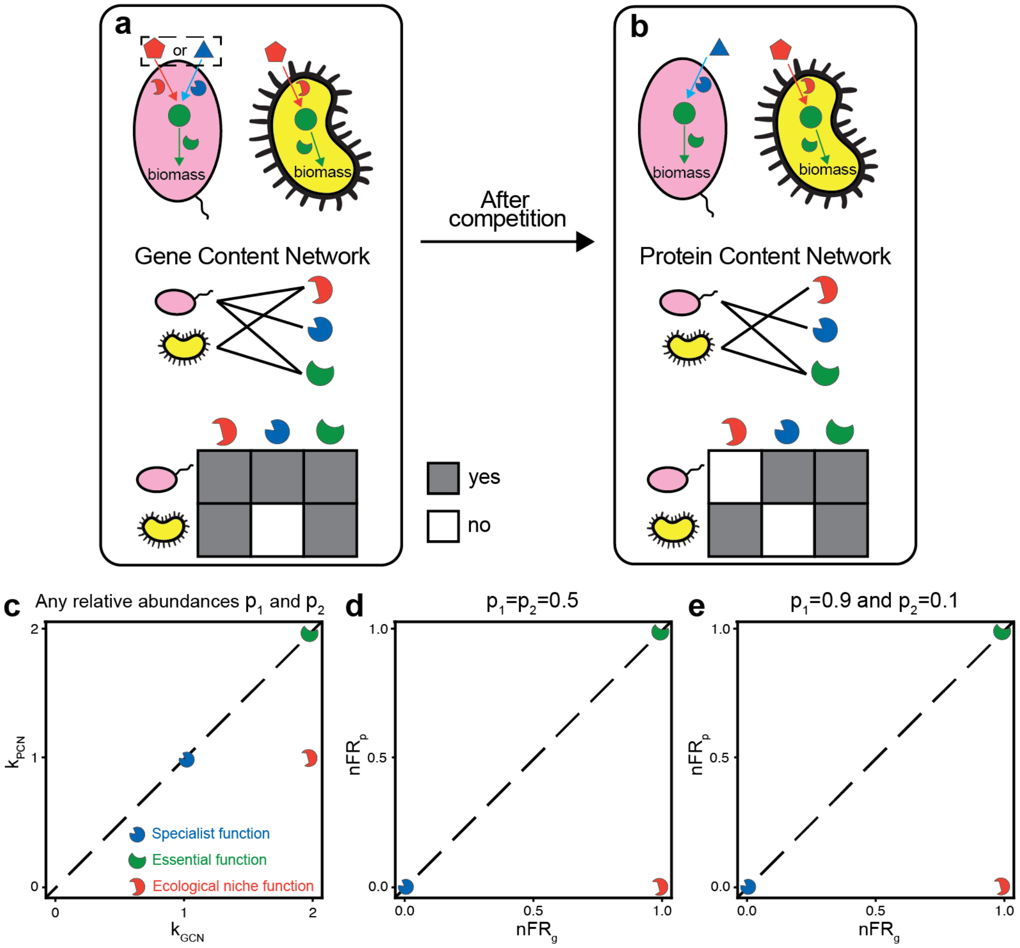


Supplementary Figure 1: **Protein functions involved in determining ecological niches are postulated to have larger discrepancies between the normalized gene-level functional redundancy $nFR_g$ and normalized protein-level functional redundancy $nFR_p$.** $nFR_g$ (or $nFR_p$) is the ratio between $FR_g$ (or $FR_p$) and taxonomic diversity TD ($= 1 - \sum_i p_i^2$). The hypothetical example used here is the same as Fig. 1.

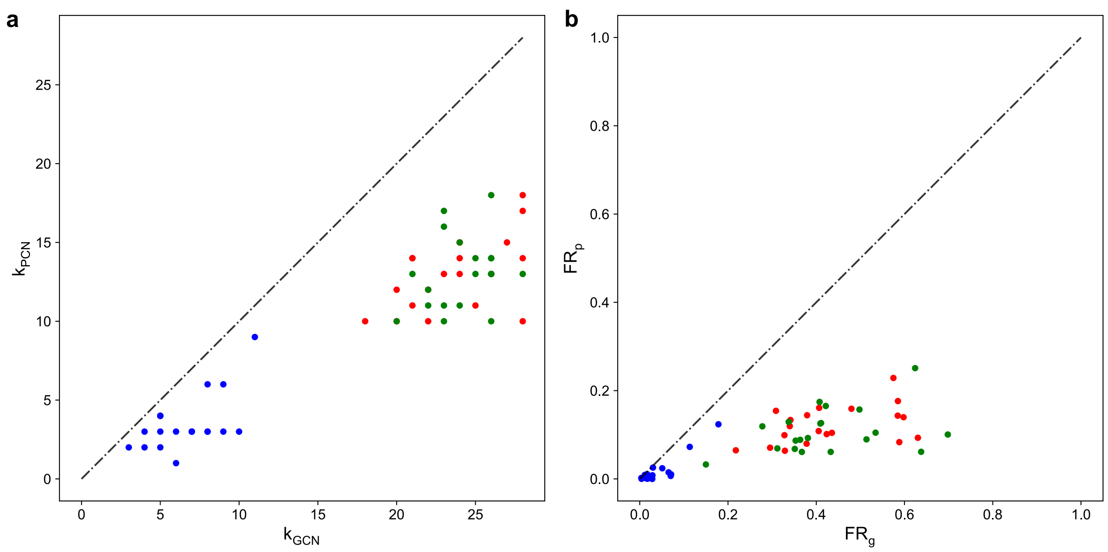


Supplementary Figure 2: **Essential functions and niche functions are indistinguishable for a randomly sampled community without going through the community assembly.** In this simulation, 20 specialist functions, 20 essential functions, and 20 ecological niche functions are modeled. 10,000 species are considered. 35 random species out of the 10,000 species in the initial pool are sampled to form a random community. The abundances of those species are assumed to be the same as the abundances of survived species in the simulation in Fig. 2c. **a-b,** The comparison of network degree and functional redundancy respectively based on the GCN and PCN of the random community. The number of drawn species is equal to the number of survived species in Fig. 2c. $FR_g$ (or $FR_p$) is the functional redundancy of each function on the gene level (or protein level). All points/functions are colored red (niche functions), green (essential functions), and blue (specialist functions) according to their types of functions in the model.

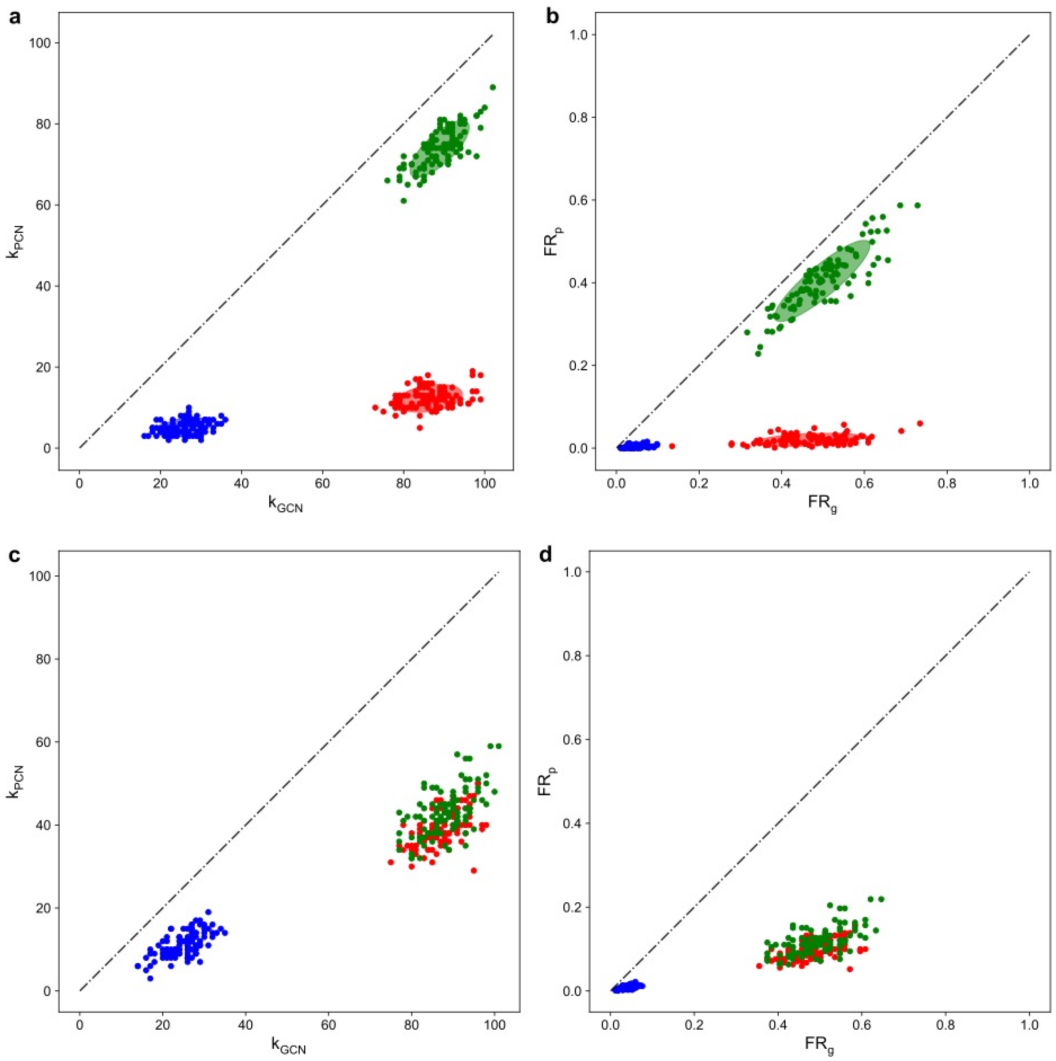


Supplementary Figure 3: **Three protein functional clusters (specialist function, essential function, and niche function) considered in the community assembly model form three distinct clusters when the network degree and functional redundancy are compared between the GCN and PCN in model-generated synthetic data.** In this simulation, 100 specialist functions, 100 essential functions, and 100 ecological niche functions are modeled. 10,000 species are considered. Eventually all species are co-cultured together to simulate their ecological competition. **a-b,** The comparison of network degree and functional redundancy respectively based on the GCN and PCN of survived species in the model simulation. kGCN (or kPCN) is the network degree of each function in the GCN (or PCN). FRg (or FRp) is the functional redundancy of each function on the gene level (or protein level). The Gaussian mixture model with 3 clusters is used to identify 3 protein functional clusters. Ellipses around clusters cover areas one standard deviation away from their means. **c-d,** The comparison of network degree and functional redundancy respectively based on the GCN and PCN of randomly drawn species in equal abundances without running community assembly. The number of drawn species is equal to the number of survived species in panels a and b. All points/functions in panels c and d are colored red (niche functions), green (essential functions), and blue (specialist functions) according to their types of functions in the model.

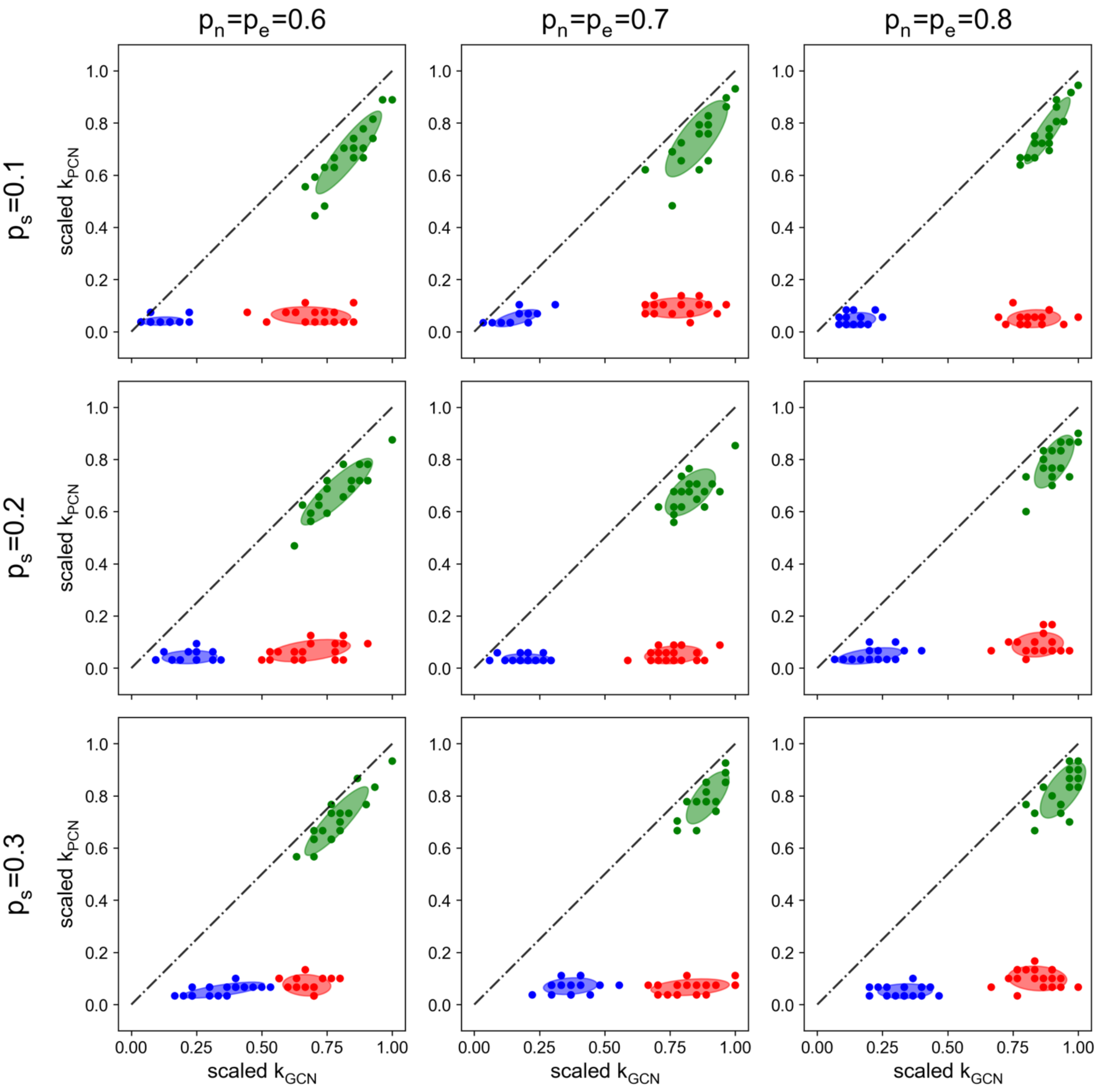


Supplementary Figure 4: **Three protein functional clusters (specialist function, essential function, and niche function) considered in the community assembly model form three distinct clusters when the network degree is compared between the GCN and PCN in model-generated synthetic data.** In this simulation, 100 specialist functions, 100 essential functions, and 100 ecological niche functions are modeled. 10,000 species are considered. Different initial probabilities of one niche (specialist, or essential) function being assigned to the species' genome, i.e., $p_n$ ($p_s$, or $p_e$), are varied. $k_{GCN}$ (or $k_{PCN}$) is the network degree of each function in the GCN (or PCN). $k_{GCN}$ and $k_{PCN}$ are linearly rescaled to maintain them between 0 and 1 so that they are visually easy to compare. The Gaussian mixture model with 3 clusters is used to identify 3 protein functional clusters. Ellipses around clusters cover areas one standard deviation away from their means. All points/functions in all panels are colored red (niche functions), green (essential functions), and blue (specialist functions) according to their types of functions in the model.

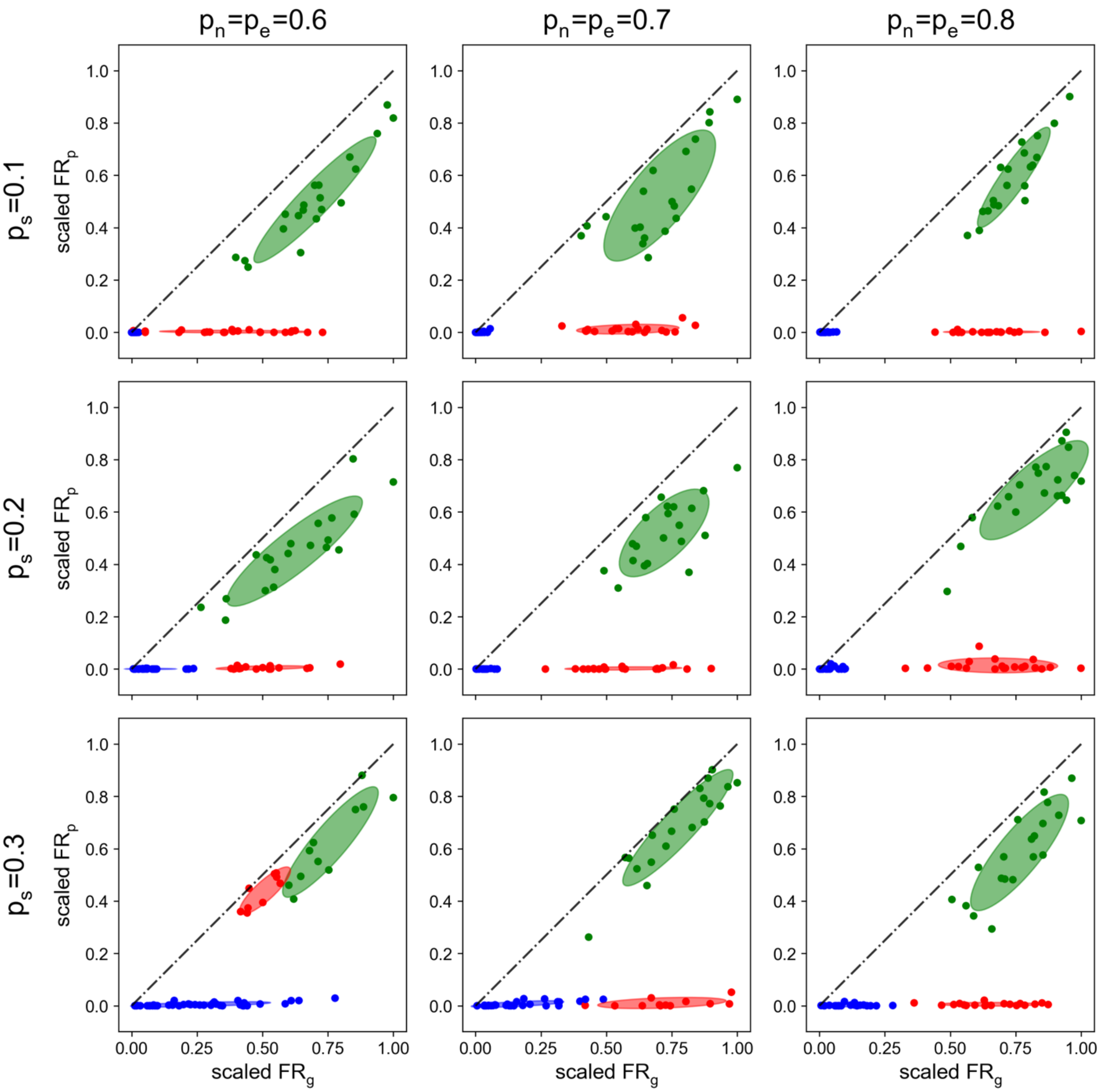


Supplementary Figure 5: **Three protein functional clusters (specialist function, essential function, and niche function) considered in the community assembly model form three distinct clusters when the functional redundancy is compared between the GCN and PCN in model-generated synthetic data.** In this simulation, 100 specialist functions, 100 essential functions, and 100 ecological niche functions are modeled. 10,000 species are considered. Different initial probabilities of one niche (specialist, or essential) function being assigned to the species' genome, i.e., $p_n$ ($p_s$, or $p_e$), are varied. $FR_g$ (or $FR_p$) is the functional redundancy of each function in the GCN (or PCN). $FR_g$ and $FR_p$ are linearly rescaled to maintain them between 0 and 1 so that they are visually easy to compare. The Gaussian mixture model with 3 clusters is used to identify 3 protein functional clusters. Ellipses around clusters cover areas one standard deviation away from their means. All points/functions in all panels are colored red (niche functions), green (essential functions), and blue (specialist functions) according to their types of functions in the model.

| | | | | | | | |
|---|---|---|---|---|---|---|---|
| *Bacteroides stercoris* | 200 | YYLFLK TTYSDIR | MVGAPPSSIGK | FGADTDNWMWPR | HTGDFSLFR | IYAGK | 236 |
| *Bacteroides fragilis* | 201 | YYLFVK TVYNDIR | MVGAPPSSIGK | FGADTDNWMWPR | HTGDFSLFR | IYADK | 237 |
| LC-MS/MS identified peptide: | | | MVGAPPSSIGK | FGADTDNWMWPR | HTGDFSLFR | | |

Supplementary Figure 6: **An illustration showing why it is difficult to construct a species-level FR based on current metaproteomic techniques.**

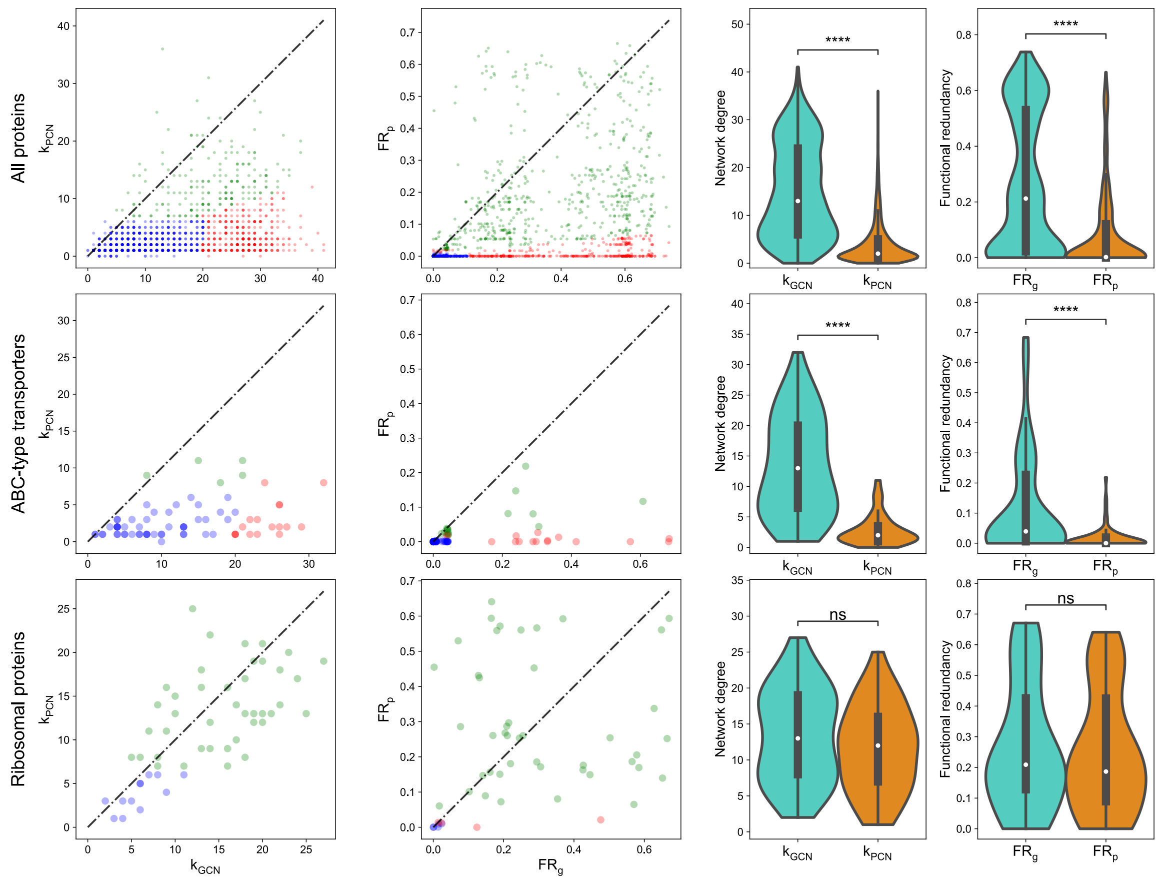


Supplementary Figure 7: **Comparison of network degree and functional redundancy between GCN and PCN annotated by KEGG as KOs (KEGG Orthologies) for the subject HM454**. We annotated inferred proteins in metagenome and metaproteome as KOs to construct the GCN and PCN respectively. $k_{GCN}$ (or $k_{PCN}$) is the network degree of each KO in the GCN (or PCN). $FR_g$ (or $FR_p$) is the functional redundancy of each KO. Three clusters with three distinct colors (blue, red, and green) are predicted by the Gaussian mixture model with 3 clusters fitted on synthetic data. The transparent large circles represent the centroids of three clusters.

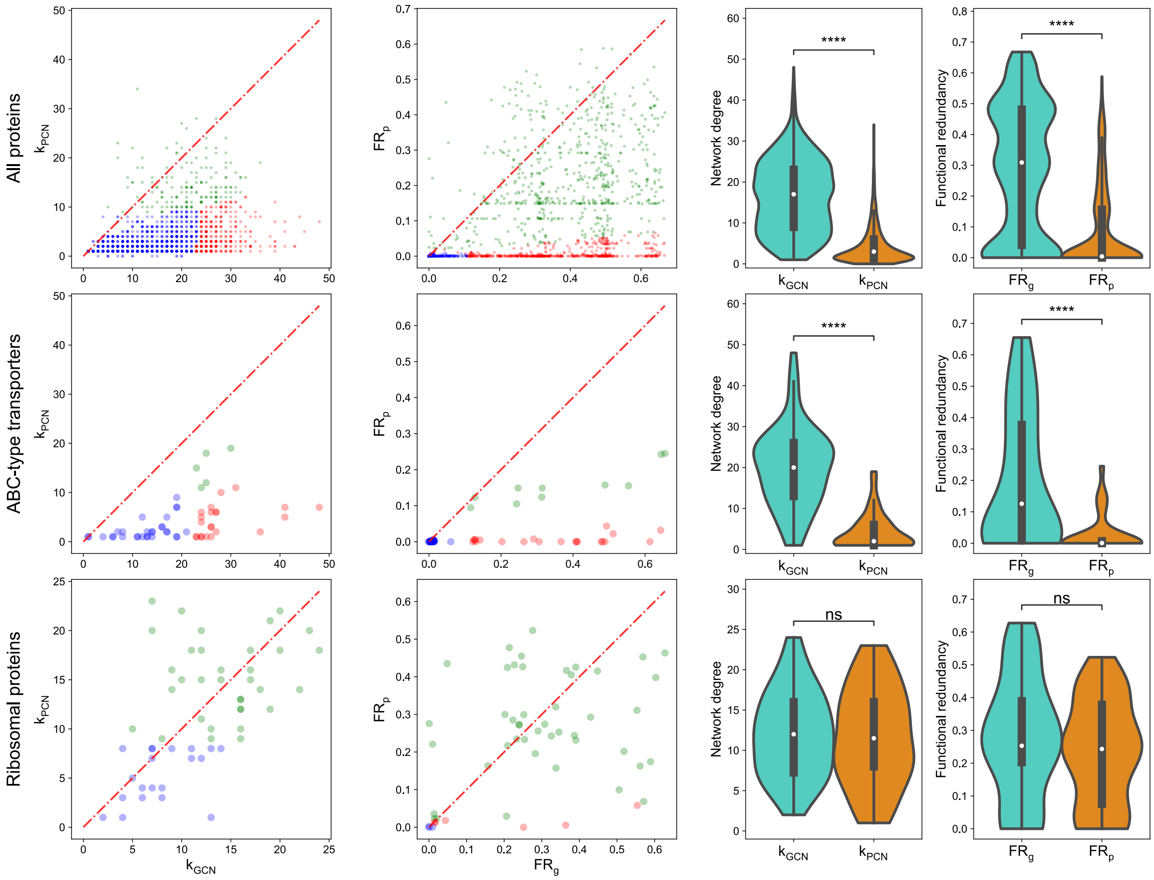


Supplementary Figure 8: **Comparison of network degree and functional redundancy between GCN and PCN annotated as COGs for the subject HM455**. $k_{GCN}$ (or $k_{PCN}$) is the network degree of each COG in the GCN (or PCN). $FR_g$ (or $FR_p$) is the functional redundancy of each COG. Three clusters with three distinct colors (blue, red, and green) are predicted by the Gaussian mixture model with 3 clusters fitted on synthetic data. The transparent large circles represent the centroids of three clusters.

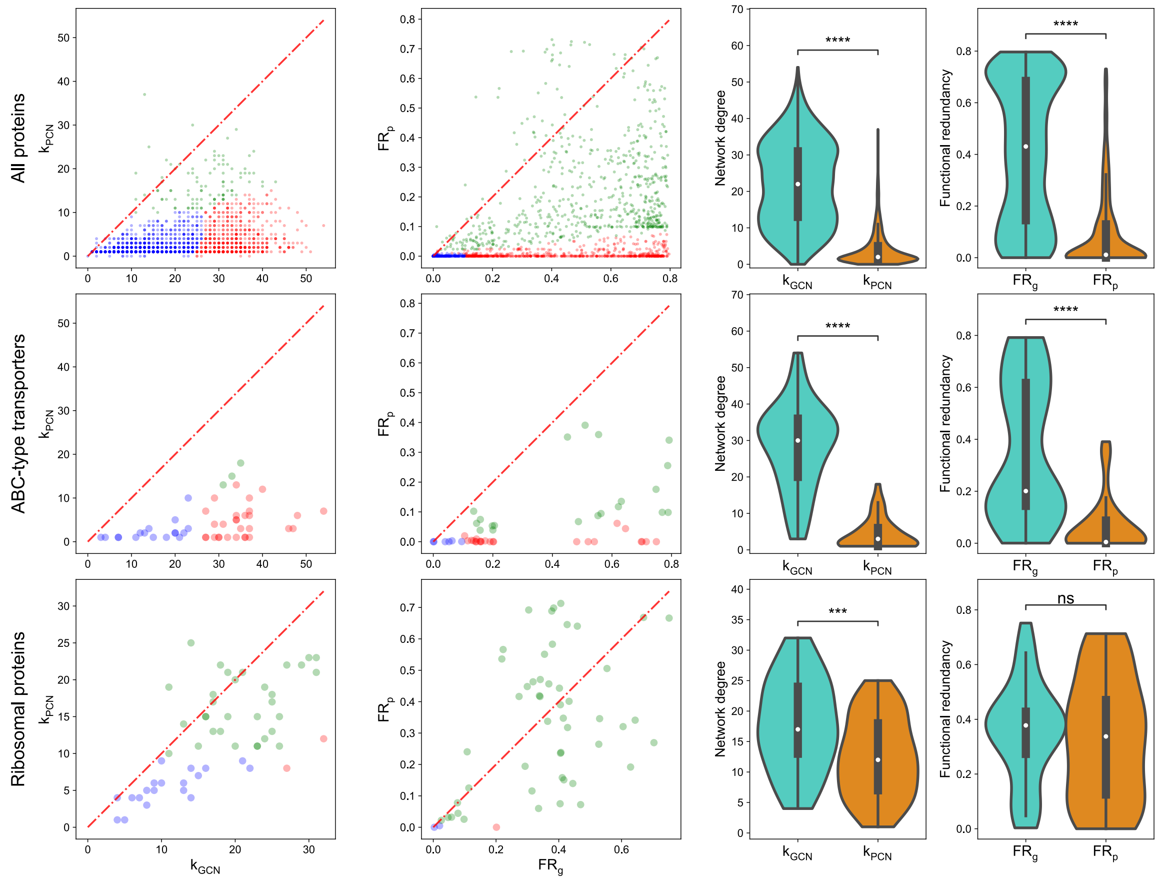


Supplementary Figure 9: **Comparison of network degree and functional redundancy between GCN and PCN annotated as COGs for the subject HM466**. $k_{GCN}$ (or $k_{PCN}$) is the network degree of each COG in the GCN (or PCN). $FR_g$ (or $FR_p$) is the functional redundancy of each COG. Three clusters with three distinct colors (blue, red, and green) are predicted by the Gaussian mixture model with 3 clusters fitted on synthetic data. The transparent large circles represent the centroids of three clusters.

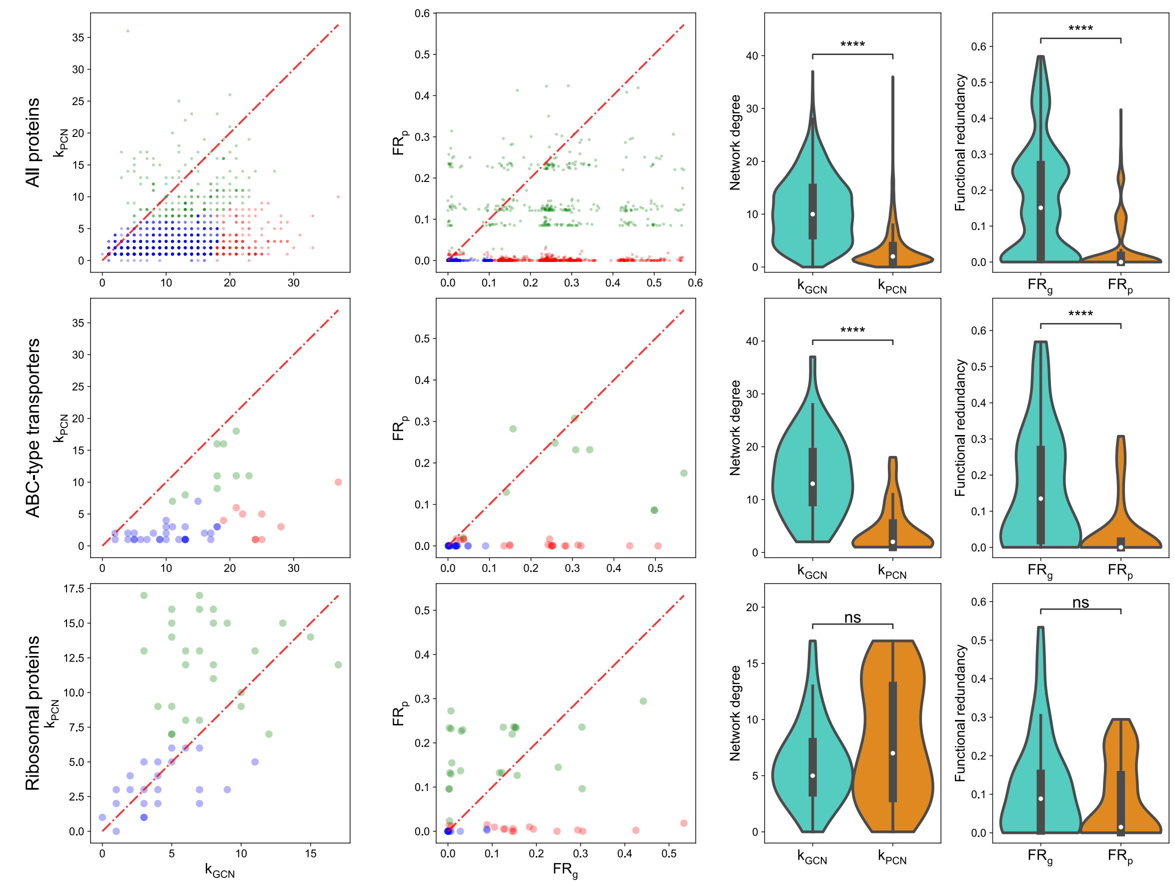


Supplementary Figure 10: **Comparison of network degree and functional redundancy between GCN and PCN annotated as COGs for the subject HM503**. $k_{GCN}$ (or $k_{PCN}$) is the network degree of each COG in the GCN (or PCN). $FR_g$ (or $FR_p$) is the functional redundancy of each COG.  Three clusters with three distinct colors (blue, red, and green) are predicted by the Gaussian mixture model with 3 clusters fitted on synthetic data. The transparent large circles represent the centroids of three clusters.

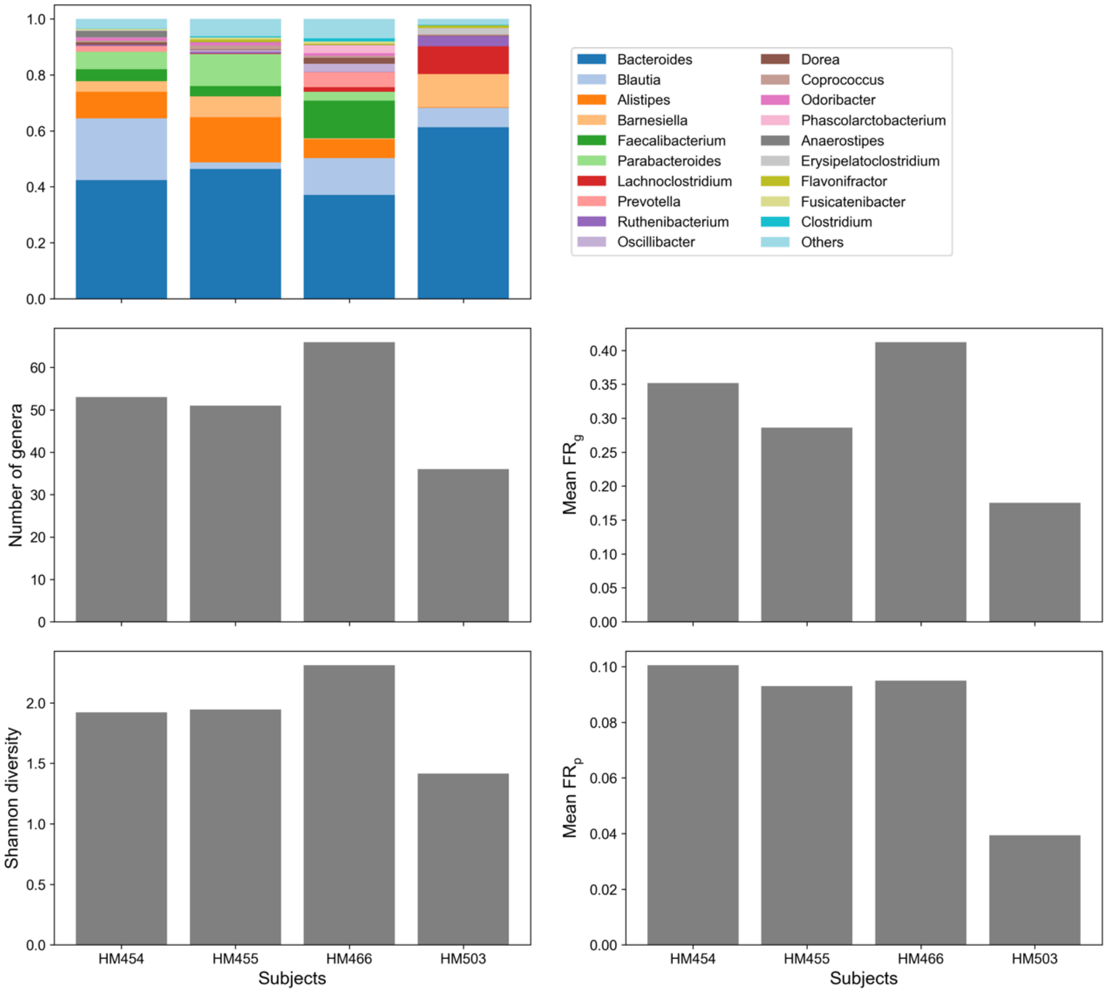


Supplementary Figure 11: **Comparison of genus composition, number of genera, Shannon diversity, mean $FR_g$, and mean $FR_p$ across the four individuals in the dataset of human gut microbiome.**

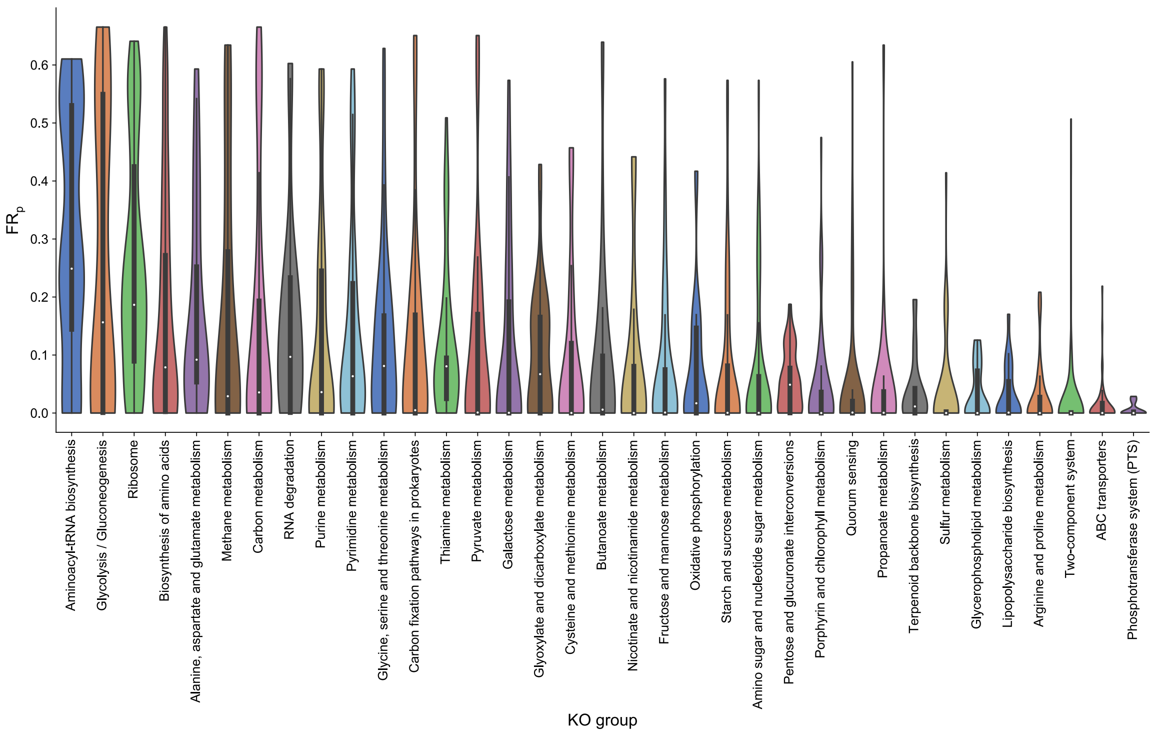


Supplementary Figure 12: **Distributions of protein-level functional redundancies $FR_p$ for different KO groups of the subject HM454.** In all boxplots, the middle white dot is the median, the lower and upper hinges correspond to the first and third quartiles, and the black line ranges from the 1.5 × IQR (where IQR is the interquartile range) below the lower hinge to 1.5 × IQR above the upper hinge. All violin plots are smoothed by a kernel density estimator and 0 is set as the lower bound.

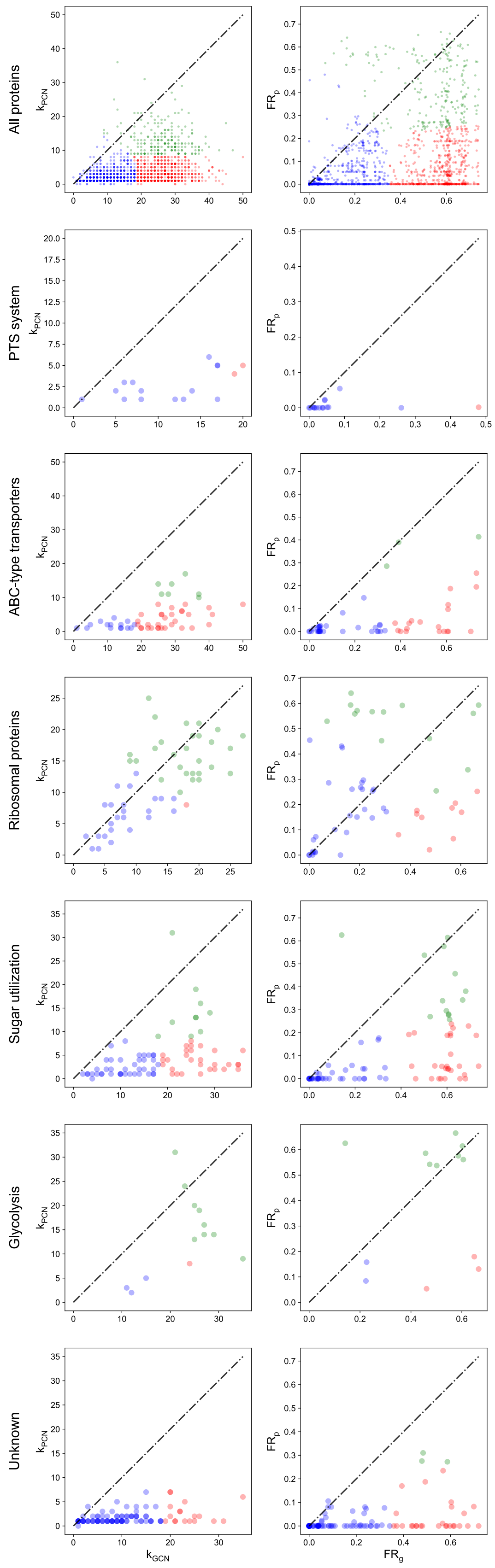
All proteins
PTS system
ABC-type transporters
Ribosomal proteins
Sugar utilization
Glycolysis
Unknown
$k_{PCN}$
$FR_p$
$k_{GCN}$
$FR_g$

Supplementary Figure 13: **K-means clustering with K=3 applied to all comparisons of network degree and functional redundancy between the gene and protein level for the human gut microbiome from HM454**. All points/functions in all panels are colored red (niche functions), green (essential functions), and blue (specialist functions) according to their types of functions in the model.

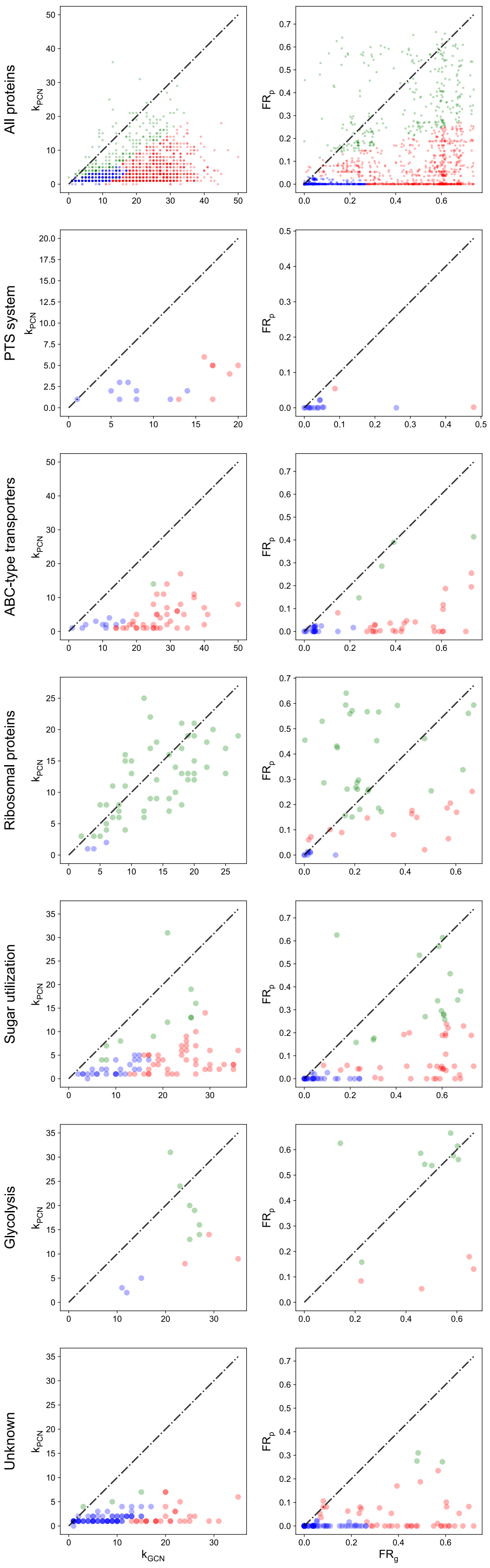

All proteins
PTS system
ABC-type transporters
Ribosomal proteins
Sugar utilization
Glycolysis
Unknown
$k_{PCN}$
$FR_p$
$k_{GCN}$
$FR_g$

Supplementary Figure 14: **Quadratic discriminant analysis (QDA) applied to all comparisons of network degree and functional redundancy between the gene and protein level for the human gut microbiome from HM454**. QDA was trained by taking the ABC-type transporters, ribosomal proteins, and PTS proteins as the niche functions, essential functions, and specialist functions respectively. All points/functions in all panels are colored red (niche functions), green (essential functions), and blue (specialist functions) according to their types of functions in the model.

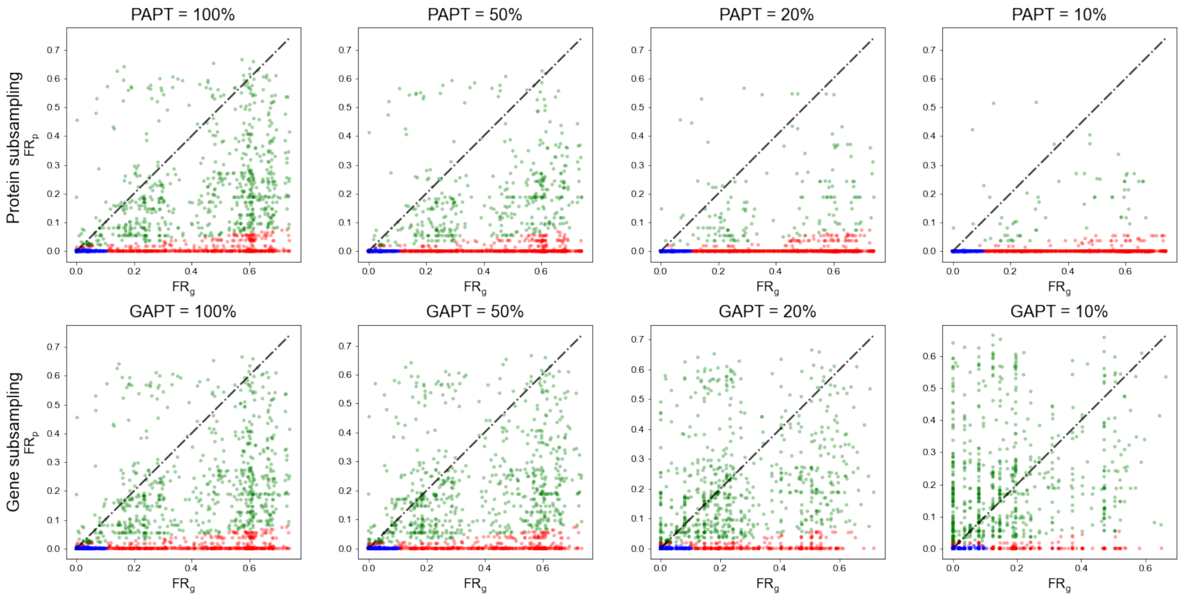


Supplementary Figure 15: **Comparison of functional redundancy between the gene and protein level ($FR_g$ and $FR_p$) for ABC-type transporters and ribosomal proteins from the human gut microbiome of HM454 when the protein or gene detection threshold changes.** Protein Abundance Percentile Threshold (PAPT) is defined as the percentage of most abundant proteins being kept. Gene Abundance Percentile Threshold (GAPT) is defined as the percentage of most abundant genes being kept.

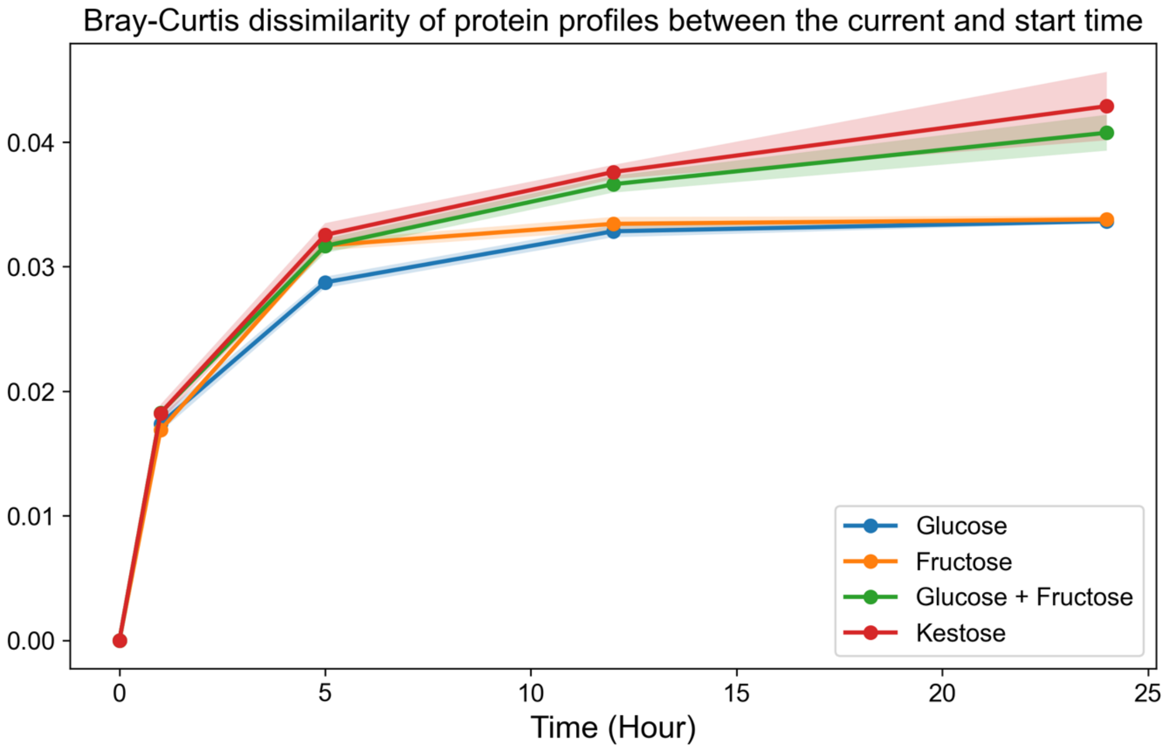


Supplementary Figure 16: **The protein profiles resulting from the introduction of more complex sugars display a greater variation compared to those observed at the initial time point.** The Bray-Curtis dissimilarity of protein profiles between the current time and the initial time is measured for four cases with different sugars introduced (glucose, fructose, glucose + fructose, and kestose).

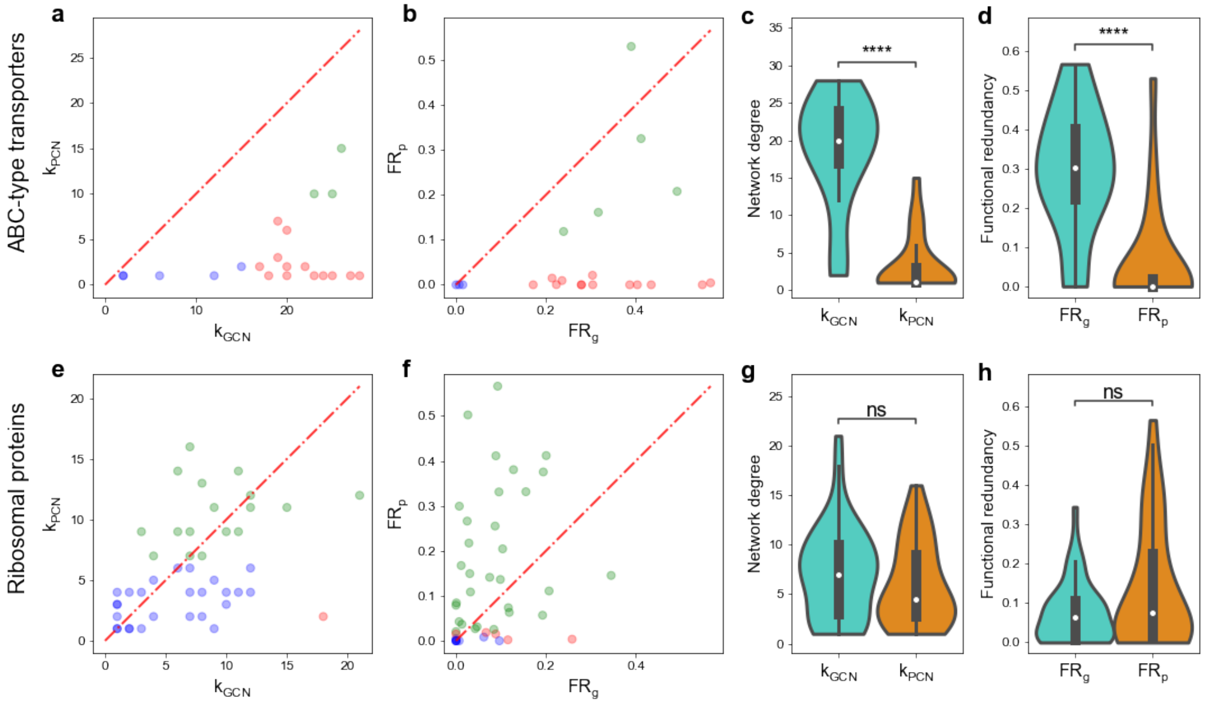


Supplementary Figure 17: **Comparison of network degree and functional redundancy between the gene and protein level for ABC-type transporters and ribosomal proteins from the *in-vitro* culture of the human gut microbiome. a,** Network degrees in GCN are larger than network degrees in PCN for most ABC-type transporter COGs. $k_{GCN}$ (or $k_{PCN}$) is the network degree of each COG in the GCN (or PCN). **b,** $FR_g$ is larger than $FR_p$ for most ABC-type transporter COGs. **c-d,** The distribution of network degrees and functional redundancies (violin plots and boxplots) for ABC-type transporter COGs show a significantly huge reduction from $k_{GCN}$ to $k_{PCN}$ or from $FR_g$ to $FR_p$. **e,** Network degrees in GCN are comparable with that in PCN for most ribosomal protein COGs. **f,** $FR_g$ is comparable with $FR_p$ for most ribosomal protein COGs. **g-h,** The distribution of network degrees and functional redundancies (violin plots and boxplots) for ribosomal protein COGs show no significant reduction from $k_{GCN}$ to $k_{PCN}$ or from $FR_g$ to $FR_p$. In all boxplots, the middle white dot is the median, the lower and upper hinges correspond to the first and third quartiles, and the black line ranges from the 1.5 × IQR (where IQR is the interquartile range) below the lower hinge to 1.5 × IQR above the upper hinge. All violin plots are smoothed by a kernel density estimator and 0 is set as the lower bound. All statistical analyses were performed using the two-sided Mann-Whitney-Wilcoxon U Test with Bonferroni correction between genomic capacity (GCN) and protein functions (PCN). P values obtained from the test is divided into 5 groups: (1) $p > 0.05$ (ns), (2) $0.01 < p \leq 0.05$ (*), (3) $10^{-3} < p \leq 0.01$ (**), (4) $10^{-4} < p \leq 10^{-3}$ (***), and (5) $p \leq 10^{-4}$ (****). Network degree comparison of ABC transporters: $p = 4.77 \times 10^{-7}$. Network degree comparison of ribosomal proteins: proteins: $p = 0.27$. Redundancy comparison of ABC transporters: $p = 1.79 \times 10^{-5}$. Redundancy comparison of ribosomal proteins: $p = 0.18$.

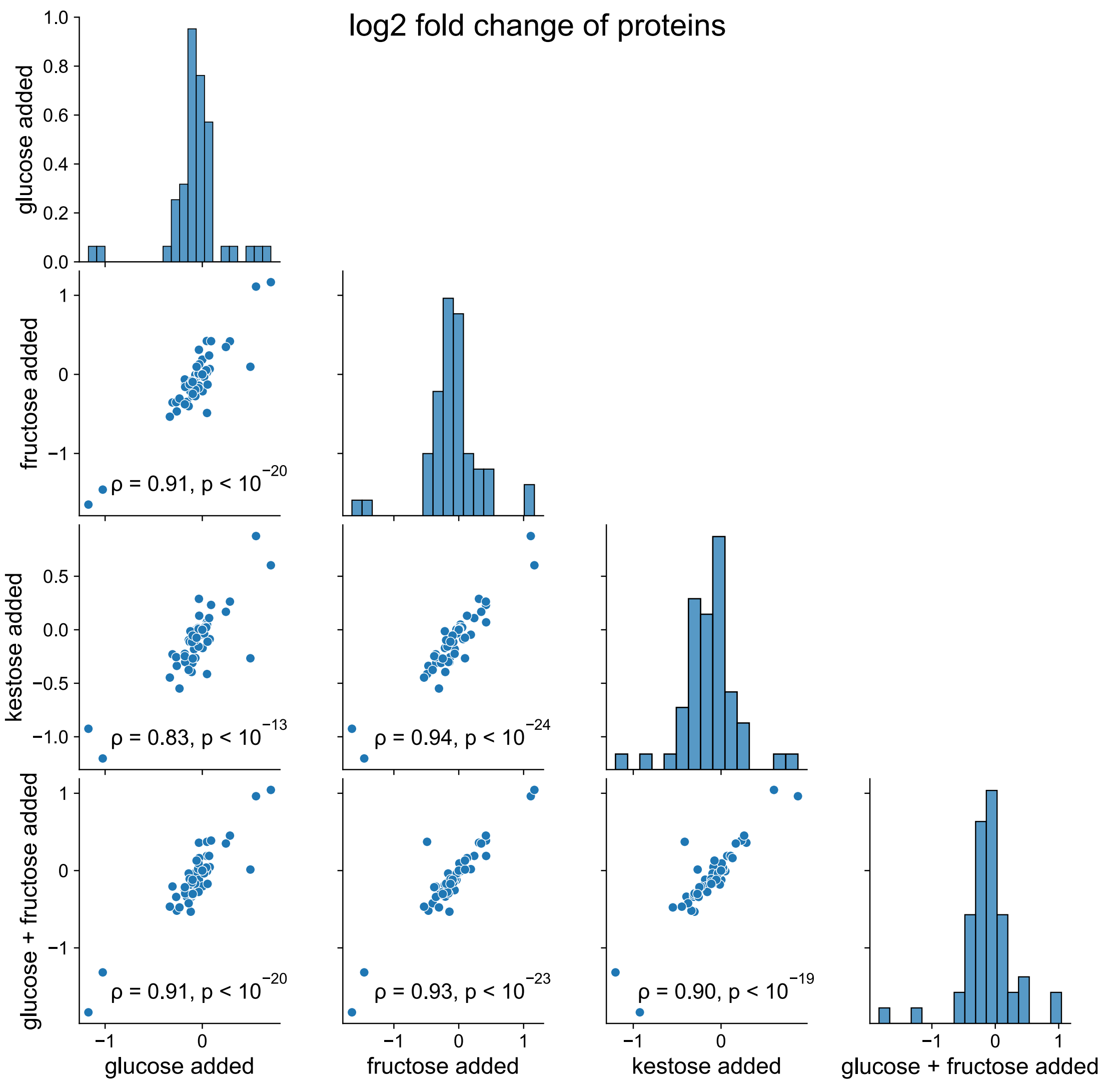


Supplementary Figure 18: **The Pearson correlation between log2 fold changes of ABC-type transporters 5 hours after different sugars are added.** The Pearson correlation coefficients ($\rho$) and corresponding p values (p) for all pairs are shown across panels.

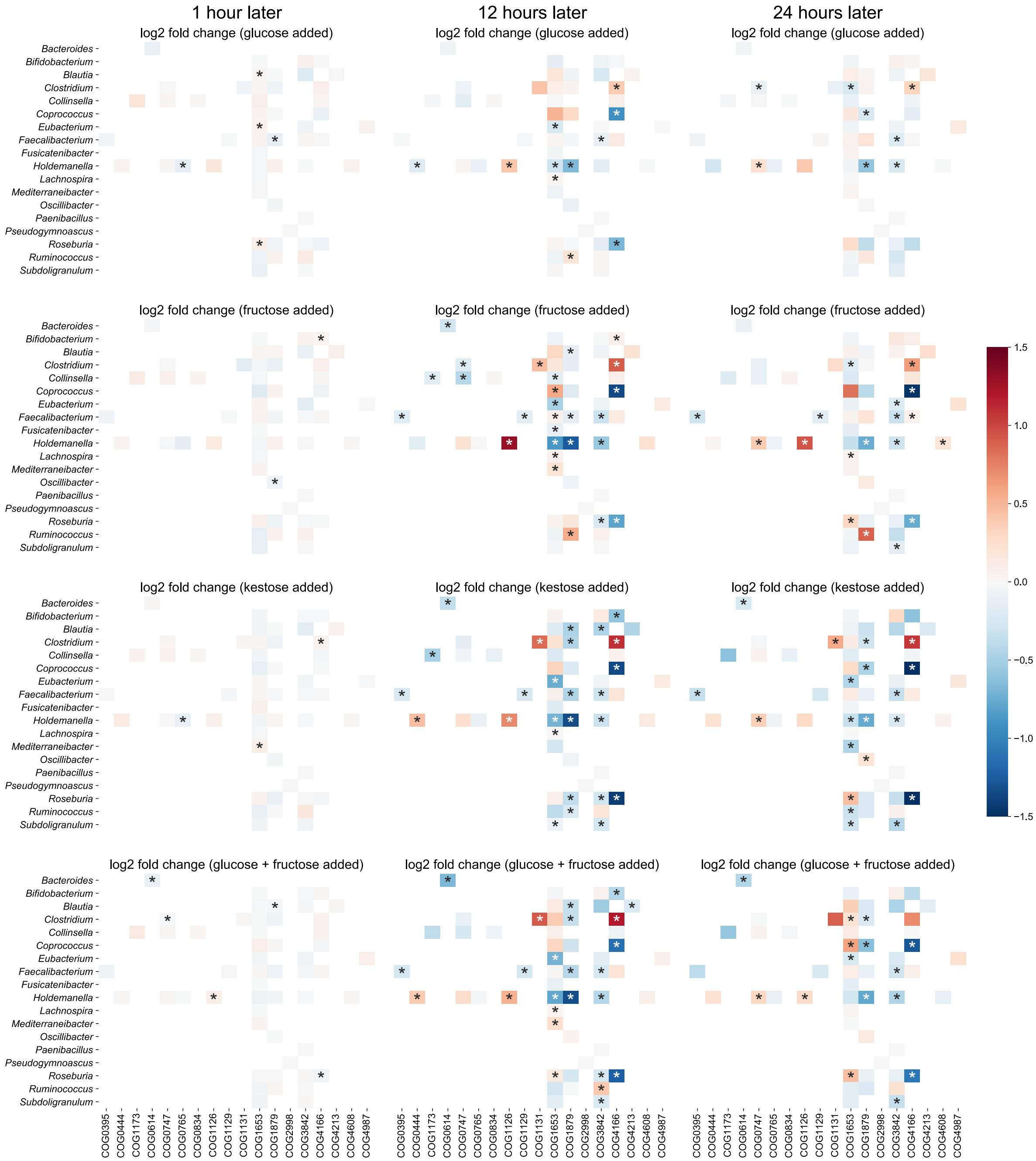


Supplementary Figure 19: **Microbes modify their expression for ABC-type transporters to adapt to different added sugars 1 hour, 12 hours, or 24 hours later.** All heatmaps share the same color bar. Metaproteomic measurements 1 hour later were used to compare the intensity of each taxon-specific protein using the log2 fold change of each protein's fraction (i.e. normalized intensity over each genus) from the treatment group divided by that from the control group.

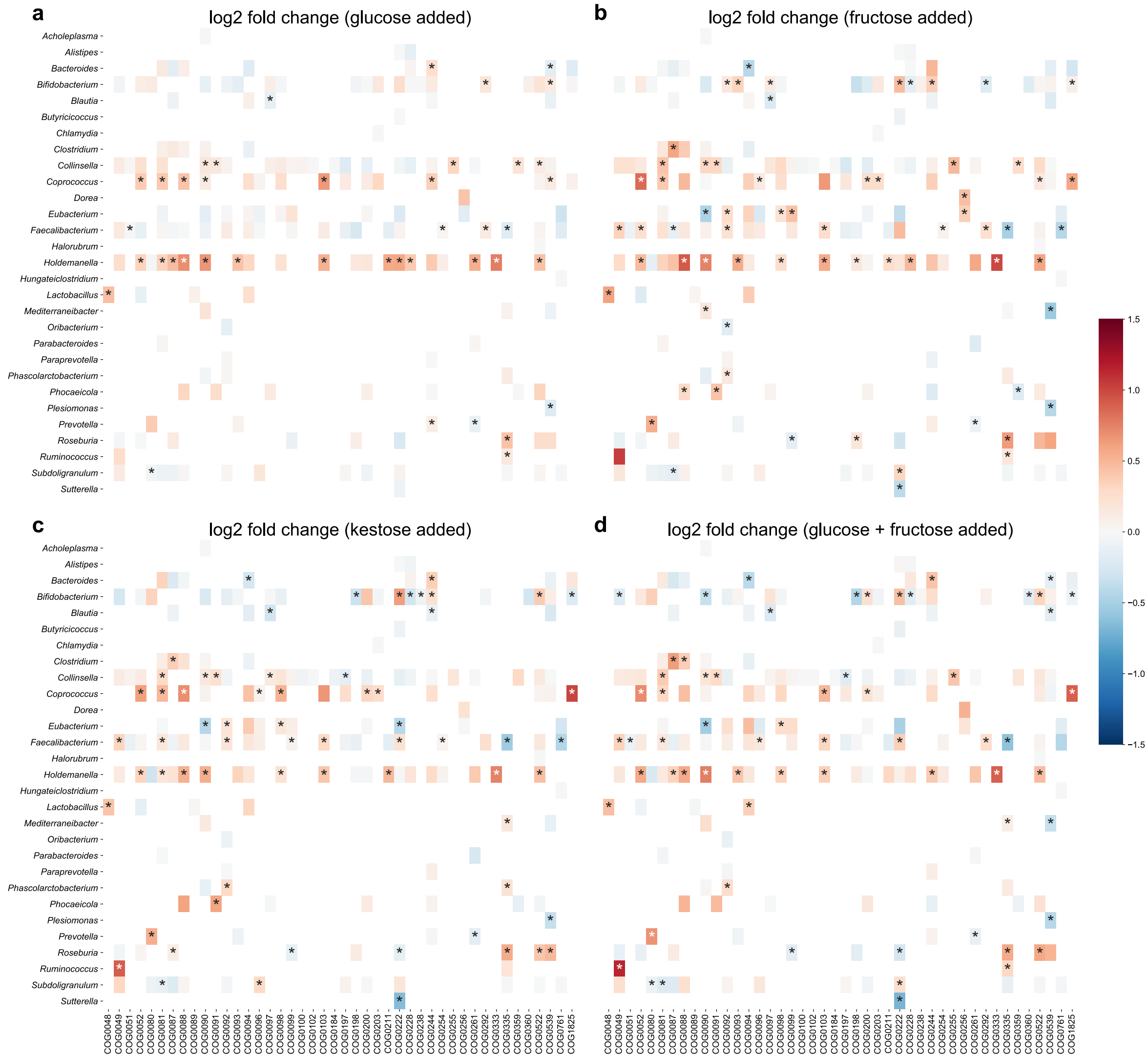


Supplementary Figure 20: **Microbes modify their expression for ribosomal proteins to adapt to added sugars 5 hours later.** All heatmaps share the same color bar shown in panel c. Log2 fold changes of ribosomal proteins were computed 5 hours after (**a**) glucose, (**b**) fructose, (**c**) kestose, or (**d**) glucose and fructose is added. We used the one-sample t-test to determine the significance of log2 fold changes and tested whether the mean of log2 fold changes from 3 experimental replicates differs from 0. Asterisks are added to positions where p values from the t-test are less than 0.05.

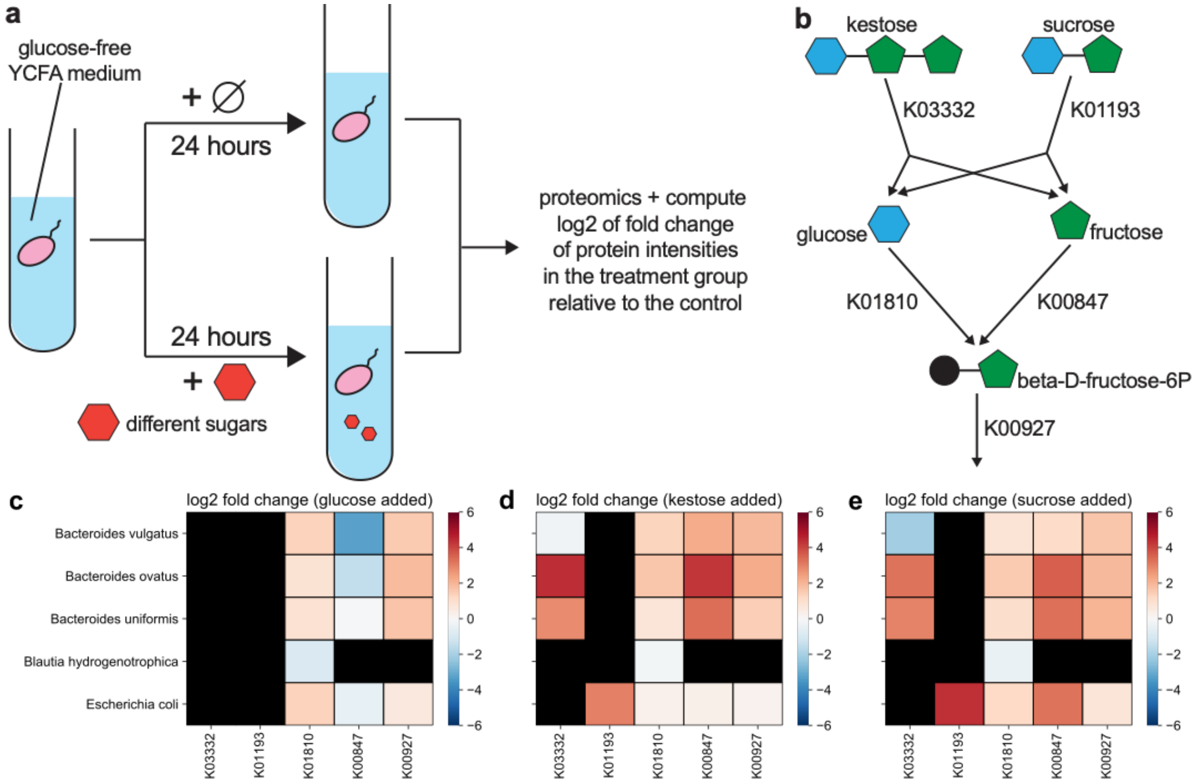


Supplementary Figure 21: **Same microbial strains have different expression levels of sugar-utilizing enzymes when different sugars are added to the glucose-free YCFA medium.** We used log2 fold changes of sugar-utilizing enzymes 24 hours after the introduction of different sugars relative to the control where no sugar is added to the base medium to reflect the difference in expression. Black colors in all heatmaps denote no measured enzymes. **a,** Schematic of *in-vitro* cultures of single microbial strains. In the treatment group, one sugar is added to the community. Metaproteomic measurements 24 hours later were used to compare the intensity of each taxon-specific protein using the log2 fold change of each protein's intensity from the treatment group divided by that from the control group. **b,** five sugar-utilizing enzymes involved in the metabolism of different sugars. Log2 fold changes of sugar-utilizing enzymes were computed 24 hours after (**c**) glucose, (**d**) kestose, or (**e**) sucrose is added.

#### Supplementary Data Legends

**Supplementary Data 1:** Table containing the network degrees (GCN and PCN), $\mathrm{FR_g}$, and $\mathrm{FR_p}$ for all annotated COGs of the subject HM454.

**Supplementary Data 2:** Table containing the network degrees (GCN and PCN), $\mathrm{FR_g}$, and $\mathrm{FR_p}$ for all annotated COGs of the subject HM455.

**Supplementary Data 3:** Table containing the network degrees (GCN and PCN), $\mathrm{FR_g}$, and $\mathrm{FR_p}$ for all annotated COGs of the subject HM466.

**Supplementary Data 4:** Table containing the network degrees (GCN and PCN), $\mathrm{FR_g}$, and $\mathrm{FR_p}$ for all annotated COGs of the subject HM503.
